# Floral evolution and pollinator diversification in *Hedychium* J.Koenig (Zingiberaceae): one of Mr. Darwin’s tropical fantasies

**DOI:** 10.1101/2021.12.06.471407

**Authors:** Ajith Ashokan, Piyakaset Suksathan, Jana Leong-Škorničková, Mark Newman, W. John Kress, Vinita Gowda

## Abstract

**PREMISE:** *Hedychium* J.Koenig (ginger lilies: Zingiberaceae) is endemic to the Indo-Malayan Realm (IMR) and is known for its fragrant flowers. Two different pollination syndromes characterize the genus: diurnal or bird pollination and nocturnal or moth pollination systems. To date, no attempt has been undertaken to understand the evolution of floral traits in this genus.

**METHODS:** We estimated ancestral character-states, phylogenetic signals, and character correlations for thirteen discrete and eight continuous floral traits representing 75% species diversity of *Hedychium*. Diversification rate estimation analyses were also employed to understand trait-dependent diversification in the genus.

**RESULTS:** Inflorescence structure, cincinnus capacity, and curvature of floral tubes revealed strong phylogenetic dependence, whereas number of open flowers per inflorescence per day, color of the labellum, and exertion of the stigma characterized higher ecological effects. Diversification rate estimations suggested that the labellum width, floral tube length, and labellum color played a major role in the evolutionary diversification of *Hedychium*.

**CONCLUSIONS:** We identified bract type and cincinnus capacity as synapomorphies for *Hedychium*, while the island-specific clade III was characterized by slender cylindrical inflorescence, coiling of floral tubes, and longer bract to calyx ratio. The circum-Himalayan clade IV is the most speciose, derived, and with most variable floral traits. Although floral color and size lacked any association with pollinator-specific traits (moth and bird pollination), pale colored flowers were most common in the early diverging clades (clade I, II-el., and II-de.), indicating their ancestral nature, when compared to brightly colored flowers.

## INTRODUCTION

***Down, March 25th [1874]***

***“I am glad to hear about the Hedychium, and how soon you have got an answer! I hope that the wings of the Sphinx will hereafter prove to be bedaubed with pollen, for the case will then prove a fine bit of prophecy from the structure of a flower to special and new means of fertilization***. *By the way, I suppose you have noticed what a grand appearance the plant makes when the green capsules open, and display the orange and crimson seeds and interior, so as to attract birds*, ***like the pale buff flowers to attract dusk-flying Lepidoptera*.*”***

**Letter 723 to J.D. Hooker**.

Flowers, a key innovation of the hyperdiverse angiosperms, are important towards our understanding of two interdependent research faculties: pollination ecology (Strauss, 1997; de Bello et al., 2021), and plant taxonomy (Sweeney, 2008; Judd et al., 2016). Ecologically, floral traits are central to the definition of pollination syndromes and pollinator interactions (Fenster et al., 2004), while taxonomically, floral characters (flowers as well as inflorescence) are important species’ descriptors. In general, many studies have shown that floral traits such as color, shape, size, fragrance, and anthesis time allow us to understand ecological interactions and their underlying selection regimes and evolutionary processes, thus emphasizing that understanding floral evolution is key to our understanding of speciation among angiosperms (Iles et al., 2017; Lagomarsino et al., 2017; Dellinger, 2020; Phillips et al., 2020).

Within the monocot order Zingiberales (8 families, 92 genera, 2,185 species; http://www.mobot.org/MOBOT/research/APweb/), the floral and inflorescence characters are known to be structurally complex (Temeles and Kress, 2003; Specht et al., 2012; Morioka et al., 2015). More than half of the species diversity in this order is concentrated within the Indo-Malayan Realm (IMR). However, our understanding of floral evolution in Zingiberales mostly stems from studies within the predominantly Neotropical families of Costaceae (Kay et al., 2005; Specht et al., 2006; Salzman et al., 2015), Heliconiaceae (Betts et al., 2015; Iles et al., 2017), Cannaceae (Specht et al., 2012), and the predominantly African groups of Marantaceae (Ley and Claßen-Bockhoff, 2009) and Strelitziaceae (Cron et al., 2012). In the IMR, despite its status as one of the eight megadiverse regions on Earth, our understanding of floral evolution among the native Zingiberales is severely lacking (Sakai et al., 1999; Paudel et al., 2019). Two factors contribute to this scarcity: a) shortages of phylogenetic frameworks of endemic lineages (Wood et al., 2000), and b) lack of systematic pollination and ecological studies in plant groups from the IMR (Specht et al., 2012).

*Hedychium* J.Koenig (ginger lilies or butterfly lilies; ∼90 species) is one of the largest genera included in Zingiberaceae (ginger family) and has a distribution exclusive to the IMR. It is the only genus in the order Zingiberales where a vast majority of species have very fragrant flowers (∼ 98% of the taxa), and this fragrance may vary throughout the day (Darwin Correspondence Project, “Letter no. 9609”; Specht et al., 2012; Ashokan and Gowda, 2019). The wide range of size and color as well as the fragrance of the flowers of *Hedychium* all contribute to the pollination syndromes in this genus (Ashokan and Gowda, 2017; Ashokan, 2020).

One of the earliest accounts in which pollination syndromes in *Hedychium* has been discussed is found in letters by Charles Darwin (Darwin Correspondence Project, “Letter no. 9372”). Towards the last quarter of the nineteenth century, in his correspondence to contemporary botanists and zoologists (Friedrich Müller, Joseph Dalton Hooker, James Alexander Gammie, and Hermann Müller), Darwin enquired about the pollination systems in *Hedychium* (Darwin Correspondence Project, “Letter no. 9371”). Upon carefully examining the floral features, Darwin predicted that the wings of moths might be able to remove and deposit pollen and result in pollination. Garden observations by F. Müller later validated Darwin’s predictions that the moths’ wings indeed transported pollen. Sir. J. D. Hooker also wrote to Darwin that the *Hedychium* flowers were moth pollinated based on field observations by J. A. Gammie (Darwin Correspondence Project, “Letter no. 9296A”). These conversations lasted for almost a decade when Darwin arranged to have samples of moths that belonged to the family Sphingidae, which were reported to have visited *Hedychium* flowers. Unfortunately, Darwin did not receive intact specimens of any of the moth samples when in London, which ended his hopes for further scientific investigations on the plant-pollinator interactions in this enigmatic group.

After more than a century from Darwin’s predictions, the only study that provided preliminary insights into the pollination syndromes in *Hedychium* was Specht et al. (2012) where a broad approach was taken in predicting the pollination syndromes across members of Zingiberales. In this study, Specht et al. (2012) identified two distinct pollination syndromes in *Hedychium*: bird pollination (ornithophily) and moth pollination (phalaenophily). However, conclusions related to the floral evolution in *Hedychium* are severely limited as Specht et al. (2012) included only 31 taxa (< 35% species diversity; all 29 taxa from the only existing molecular phylogenetic reconstruction by Wood et al., 2000) in their phylogenetic analysis. In addition to the reduced representation of taxa in both Wood et al. (2000) and Specht et al. (2012), we also identified two other causes - (i) inadequately documented floral trait data and hypothetical trait assignments, and (ii) problems in taxonomic delimitations presented by species complexes, persisting mainly as natural hybrids and polyploids (Ashokan, 2020; Ashokan et al., in prep.). When species cannot be distinguished based on morphological differences, it gives rise to the problem of species complexes (Grant, 1971; Specht, 2006; Surveswaran et al., 2018; Saryan et al., 2020). Within *Hedychium*, multiple such complexes have been identified, including the one in which the type species, *H. coronarium* J.Koenig, is also involved (Turrill, 1914; Ashokan, 2020; Saryan et al., 2020), and thus pose serious difficulties in the taxonomic identification of lineages (Saryan et al., 2020; Ashokan, 2020).

Here we build one of the most comprehensive phylogenetic trees of *Hedychium* through multi-institutional collaborations that allowed us to sample most of the taxonomic diversity of *Hedychium* and supplement floral trait information across the genus. We rephrase the two syndromes (bird and moth pollination) identified in *Hedychium* as diurnal and nocturnal pollination syndromes, respectively. This rephrasing allows us to focus on floral traits as pollination syndromes without being biased about the type of pollinators associated with the syndrome. Broadly, we divided our research aims into three major arenas: (i) define synapomorphies of clades in the genus to test all earlier subgeneric classifications of *Hedychium*, which have been based only on a few floral characters (see Ashokan and Gowda (2019) for a detailed review), (ii) estimate the evolutionary transitions in floral traits by examining pollination syndromes from two non-exclusive axes: solitary (individual flowers) and collective (whole inflorescence) floral traits, and (iii) determine the roles of these traits in the diversification of this genus. We anticipate that variation in floral traits often corresponds to allied interspecific attributes in the case of morphological intermediates (hybrids and/or polyploids), and hence an understanding of the evolution of floral traits is important to apply them as taxonomic delimitators. Using an ultrametric tree with a suite of categorical and continuous floral or inflorescence characters, we address the following questions: (1) Which floral traits define the clades within *Hedychium*? (2) Have nocturnal flowers evolved only a single time in *Hedychium*? (3) Do floral trait diversification correspond to the lineage diversification in *Hedychium*?

## MATERIALS AND METHODS

### Phylogenetic reconstruction

In order to obtain ultrametric trees suitable for comparative methods, we reconstructed a dated tree of *Hedychium* inferring evidence from four markers (ITS, *matK/trnK, rps16*, and *trnL*) that were additionally sampled for selected taxa across all the eight zingiberalean families (Ashokan et al. in review; Appendix S1a).

Bayesian relaxed molecular clock model with uncorrelated lognormal rates was specified in BEAST version 1.8.2 (Drummond et al., 2012). The birth-death process prior was selected against the Yule process prior after comparing their Bayes factors based on Maximum Likelihood Estimations (MLEs) of preliminary runs as implemented in Tracer version 1.6. Three fossils and a secondary calibration were applied in this study based on our previous analyses (Ashokan et al. in review). We conducted five independent MCMC runs of 100 million generations with sampling every 10000 generations. We combined the five independent runs with LogCombiner version 1.8.2, setting the burn-in to 25% of the initial samples of each run. We assessed the convergence with the R package ‘rwty’ version 1.0.2 (Warren et al., 2017) and checked the effective sample sizes (ESS > 200) in Tracer version 1.6.

Further, we used TreeAnnotator version 1.8.2 to obtain the maximum clade credibility (MCC) tree for all downstream comparative phylogenetic methods which are discussed henceforth.

### Morphological character coding

We exhaustively examined all *Hedychium* protologues, including monographs and revisions (see Ashokan and Gowda, 2019). Herbarium collections, including type specimens, were consulted by the first author at ARUN, ASSAM, BHPL, BM, BO, BSA, BSHC, BSI, CAL, E, K, LINN, LIV, MH, QBG, SING, and TBGT (refer to Thiers, 2016), and in online databases (Chinese Virtual Herbarium: http://www.cvh.ac.cn/en; Global Plants: https://plants.jstor.org/; Herbarium WU: https://herbarium.univie.ac.at/database/search.php; Muséum national d’Histoire naturelle: https://science.mnhn.fr/; Smithsonian Institution: https://www.si.edu/; and, Zingiberaceae Resource Centre: http://padme.rbge.org.uk/ZRC/). Metadata collection included scoring of morphological, phenological as well as ecological characters from the field, botanic gardens, and herbaria. Characters were chosen according to our prior knowledge of their taxonomic and ecological importance, and personal observations as part of our ongoing ecological and phylogeographic studies in *Hedychium* (see Appendix S1a for the characters and character states). We scored 21 floral (13 categorical and eight continuous) characters across 70 taxa of *Hedychium*, and a character matrix is constructed using MESQUITE version 3.31 (Maddison and Maddison, 2017).

### Ancestral character-state reconstruction and phylomorphospace analysis

All 13 categorical (binary and multistate) characters were mapped onto the phylogenetic tree using both likelihood and stochastic methods (Bollback, 2006; Maddison and Maddison, 2017). We first used the “ace” function in the R package ‘ape’ version 5.3 (Paradis et al., 2004) to estimate the likelihood of the ancestral state using a one-parameter equal-rates model (ER), three-parameter symmetrical model (SYM), and six-parameter all-rates-different model (ARD). We then used the “make.simmap” function in the R package ‘phytools’ version 0.7-70 (Revell, 2012) to estimate ancestral character states to identify putative synapomorphies in *Hedychium* (morphological characters targeted: slender cylindrical inflorescence, inflorescence appearance, inflorescence rachis exposure, inflorescence bract type, cincinnus capacity, floral tube coiling, relative length of bract and calyx, type of labellum claw, type of stigma) and detect evolutionary transitions (morphological characters targeted: floral density, number of flowers open per inflorescence per day, the color of the labellum, color of labellum blotch). We simulated 1000 mappings to approximate the posterior distribution. We then compared the fit of these models to the data by comparing the AICc score. Finally, the posterior probabilities from stochastic mapping were compared with the marginal ancestral states from the likelihood estimation. Based on these analyses we (i) reconstructed the evolution of floral characters in *Hedychium*; and (ii) tested whether the evolution of floral characters in *Hedychium* followed the patterns predicted by the floral syndromes (Grant, 1950; Specht et al., 2012).

We also assessed the disparity through time for the eight continuous characters using the function, “dtt” in the R package ‘geiger’ version 2.0.7 (Harmon et al., 2020). Further, we performed an NMDS analysis for the 13 categorical and eight continuous variables using the “metaMDS’’ function in the R package ‘vegan’ version 2.5-7 (Oksanen et al., 2013). In order to estimate the phylomorphospace in *Hedychium* we mapped NMDS1, NMDS2, and NMDS3 to the ultrametric tree, using the function, “phylomorphospace” in the R package ‘phytools’ version 0.7-70 (Revell, 2012).

### Phylogenetic signal and character correlations among categorical characters

Character correlations were estimated for different combinations of characters that were discussed in Appendix S1a. We assessed the character correlation using the “fitPagel” function in R package ‘phytools’ version 0.7-70 (Revell, 2012). Both ER and ARD models were employed for all the characters, after specifying the “fitDiscrete” method. Finally, the choice of independent or dependent (or correlated) models were determined using their AICc values and the correlations were ascertained using the *P*-value (< 0.05).

Phylogenetic signals for the 13 categorical characters that were employed in the ancestral character-state reconstruction were estimated using the “phylo.d” function in the R package ‘caper’ version 1.0.1 (Orme et al., 2013). We transformed the multistate discrete traits (color of labellum) into binary discrete traits (Appendix S1). The D statistic is negatively correlated with Pagel’s λ as the latter estimates the correlation of continuous traits from 0 to 1 (0 indicates no phylogenetic dependence and 1 indicates strong phylogenetic dependence). The D value of ‘1’ corresponds to a phylogenetically random trait and ‘0’ for a trait evolved under the Brownian model or evidence of strong phylogenetic dependence (Fritz and Purvis, 2010). The values of D above ‘1’ correspond to higher ecological effects and values below ‘0’ indicate stronger phylogenetic dependence. Recently, Borges et al. (2019) warned against using the D statistic where categorical traits do not evolve according to a Brownian motion threshold model. Hence, we also estimated the phylogenetic signal for the 13 categorical characters using the δ-statistic (Borges et al., 2019) applying the “delta” function in the R package ‘ape’ version 5.3 (Paradis et al., 2004). The δ can be any positive real number and the higher the δ-value, the higher the degree of phylogenetic signal between a given trait and the phylogeny. If the associated *P*-value is less than the level of the test (generally 0.05), it represents the presence of a phylogenetic signal between the trait and the tree. In contrast, if the *P*-value is greater than the level of test (generally 0.05) it represents lack of evidence for phylogenetic signals or that the trait is saturated.

### Phylogenetic signal and character correlations among continuous characters

To map all eight continuous characters in *Hedychium* we used the “contMap” function in the R package ‘phytools’ version 0.7-70 (Revell, 2012). Both Pagel’s λ (Pagel, 1997; Pagel, 1999) and Blomberg’s K (Blomberg and Garland, 2002) were estimated for the eight continuous traits (Appendix S1d) using the function “phylosig” in the R package ‘phytools’ version 0.7-70 (Revell, 2012) to test the hypothesis of phylogenetic signal variation among floral traits in *Hedychium*. Four different evolutionary models—Brownian motion model (BM), Ornstein-Uhlenbeck model (OU), Early Burst (EB), and White noise model (WN)—were assessed with the function “fitContinuous” in the R package ‘geiger’ version 2.0.7 (Harmon et al., 2020) to determine which model best explains the evolutionary patterns of the traits. The classic BM evolutionary model assumes random walk divergences in species resemblance (Felsenstein, 1973). The OU model is a modification of the BM model with an additional parameter, alpha (α) or ’’rubber band’’ parameter, which measures the strength of return towards a theoretical optimum that is shared across a clade or subset of species (Hansen, 1997). The EB model assumes a sudden burst in the evolution of characters in the initial stages and further normalization with time (Harmon et al., 2010). The WN model assumes no phylogenetic signal and that the data come from a single normal distribution with no covariance structure among species (Harmon et al., 2010).

Correlations among labellum, floral tube, and filament traits were expected because they are potentially closely linked to pollination. We specifically focused on four variables—the labellum length, labellum width, floral tube length, and filament length—which were indicated as having contrasting evolutionary patterns based on the estimated Pagel’s λ and Blomberg’s K values. To test whether these contrasting patterns can be predicted by the other four putatively associated traits (the predictor variables), we regressed labellum length, labellum width, floral tube length, and filament length (the response variables) against the values of four individual traits and a combination of four other traits with phylogenetic generalized least squares (PGLS; Martins and Hansen, 1997), using the function “pgls” in the R package ‘caper’ version 1.0.1 (Orme et al., 2013).

### Trait dependent diversification analyses

To estimate if there is any trait-dependent diversification within *Hedychium*, we used all 13 categorical characters independently, using the R package ‘hisse’ version 1.9.10 (Beaulieu and O’Meara, 2016). HiSSE is a state-dependent speciation and extinction form of a hidden Markov model and it assumes that related to each observed state in the model are certain “hidden” states that display potentially distinct diversification dynamics and transition rates than the observed states alone (Beaulieu and O’Meara, 2016). Of the thirty different models tested, twelve were binary state speciation and extinction (BiSSE)-like models that excluded hidden states or constrained specific parameters of turnover, extinction, or transition rates, another twelve corresponded to hidden state-dependent models with various constraints on the above parameters among states, and six were null HiSSE models with various character-independent diversification (CID) forms.

We estimated the trait-dependent diversification or phenotype evolution for the eight continuous traits using the program BAMM version 2.5.0 (Bayesian Analysis of Macroevolutionary Mixtures; Rabosky, 2014; Rabosky et al., 2017). The BAMM analysis was conducted with four reversible jump MCMC runs for 100 million generations, sampling every 10000 generations. The prior distributions were estimated with the “setBAMMPriors” function in the R package ‘BAMMtools’ version 2.1.6 (Rabosky et al., 2017). MCMC convergence was assessed in the R package ‘coda’ version 0.19-2 (Plummer et al., 2006) by checking the ESS values for likelihood and number of shift events. The first 10% of the sampled generations were discarded as burn-in and ESS values above 200 were considered indicative of convergence. The dynamic rate variation among lineages was evaluated using the following approaches in BAMMtools: (i) mean phylorate plot displayed distinct diversification rates by mapping colors to rates on all branches; and (ii) macroevolutionary cohort matrix displayed the pairwise probability that any two species shared a common macroevolutionary rate dynamic.

## RESULTS

### Phylogenetic reconstruction from the combined molecular datasets

The topology resulting from the BEAST MCMC analyses recovered five clades (clade I or Spicatum clade, clade II-el. or Ellipticum clade, clade II-de. or Densiflorum clade, clade III or Villosum clade, clade IV or Coronarium clade) and *H. pauciflorum* as the early diverging species (Ashokan et al., in review.).

Cincinnus capacity (Character 6; Fig. 1A) was identified as the only character distinguishing different clades of *Hedychium*. Clades I, II-el., and II-de. consisted of taxa with one or two flowers per bract whereas clades III and IV were made up of taxa with multi-flowered cincinni. Both slender cylindrical inflorescence (Character 1; Fig. 1C) and the coiling of floral tubes (Character 9; Fig. 1D) were identified as synapomorphies for clade III.

**FIGURE 1.**
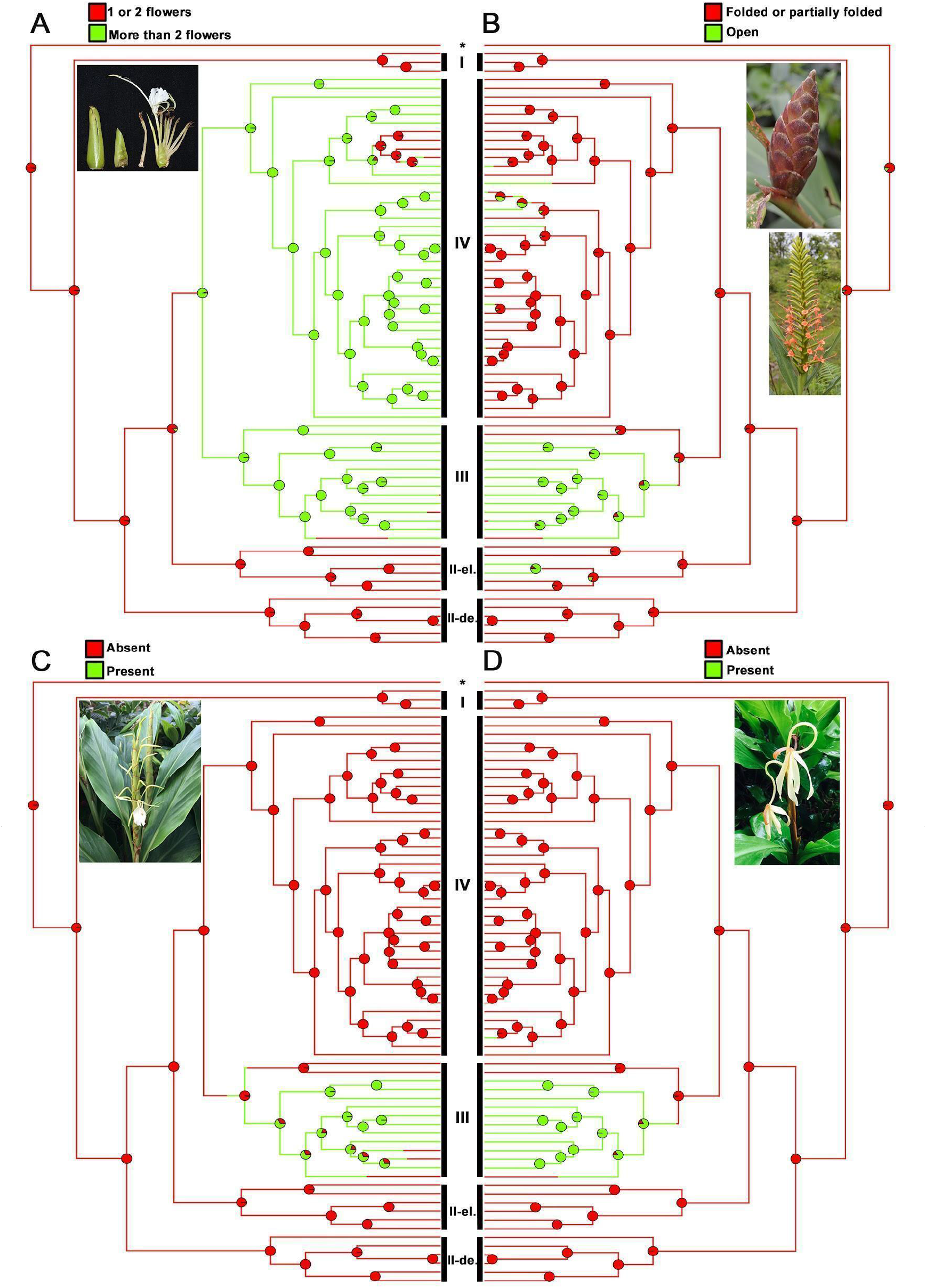
Ancestral character-state reconstructions of discrete floral traits in *Hedychium*. (A) Cincinnus capacity, (B) Inflorescence bract type, (C) Slender cylindrical inflorescence, and (D) Coiling of the floral tube.

### Ancestral character-state reconstructions

Of the 13 categorical characters, seven are inflorescence specific, and six are specific to flowers. All seven inflorescence characters (shape, appearance, rachis exposure, bract type, cincinnus capacity, floral density, number of flowers open per inflorescence per day) and four floral characters (floral tube coiling, ratio of relative lengths of bract and calyx, labellum claw type, and labellum blotch color) were supported by the ER model. Only two floral characters: the color of labellum and stigma exsertion, were best described by the ARD model. Stochastic ancestral character-state reconstructions of the 13 discrete characters are illustrated in Fig. 1, Fig. 2, and Appendix S2a. Appendix S1a presents the comparison of all three models used in the ancestral character-state reconstructions.

**FIGURE 2.**
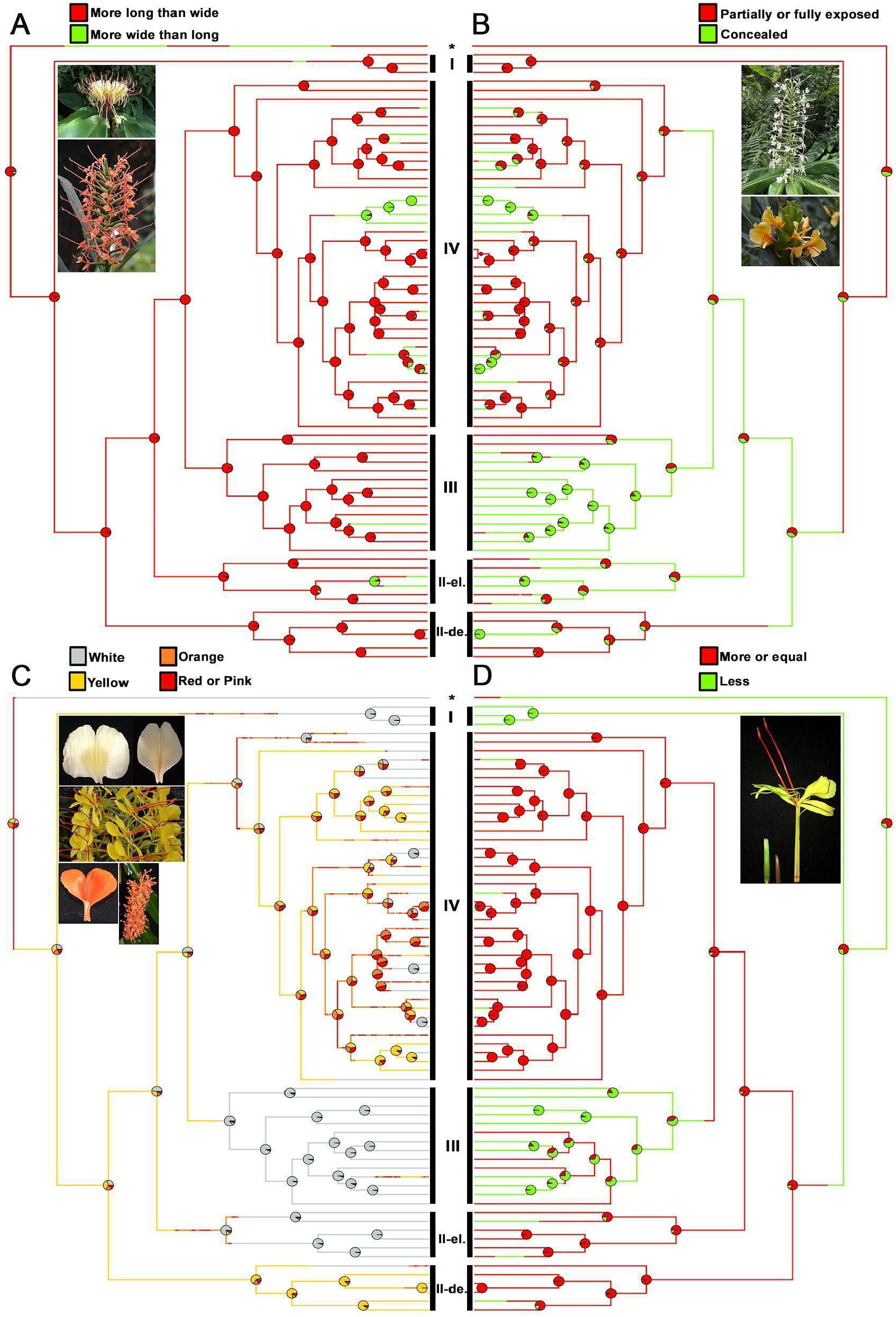
Ancestral character-state reconstructions of discrete floral traits in *Hedychium*. (A) Appearance of inflorescence, (B) Exposure of inflorescence rachis, (C) Color of labellum (excluding blotch), and (D) Ratio of relative lengths of bract and calyx.

From the ancestral character-state reconstructions, the following traits were identified as plesiomorphic for the genus *Hedychium*: inflorescence bracts with one or two flowers (Character 6; Fig. 1A), folded bract type (Character 4; Fig. 1B), absence of slender cylindrical inflorescence (Character 1; Fig. 1C), absence of floral tube coiling (Character 9; Fig. 1D), longer than wide inflorescence (Character 2; Fig. 2A), partially or wholly exposed rachis (Character 3; Fig. 2B), longer bract with respect to the calyx (Character 8; Fig. 2D), and dense or moderately dense inflorescence (Character 5; Appendix S2aA).

The open bract type evolved at least three times independently in the phylogeny of *Hedychium* (clades I, II-el., and IV). The number of flowers opens per inflorescence per day (Character 9; Appendix S2aB), type of labellum claw (Character 9; Appendix S2aC), labellum blotch color (Character 9; Appendix S2aD) and exsertion of stigma (Character 9; Appendix S2aE) are also found to have evolved multiple times independently in the phylogeny of *Hedychium*. Importantly, both white and non-white flowers evolved multiple times in the phylogeny of *Hedychium* (Character 11; Fig. 2C), with all floral colors of *Hedychium* represented only in clade IV.

### Character correlations and phylogenetic signal among categorical characters

Of 78 total correlation analyses performed, 28 of them were found to be correlated (Appendix S1b and S1c). Among the solitary floral characters, the color of the labellum showed the most number of correlations whereas, among the collective floral characters (or inflorescence characters), the inflorescence bract type has the most number of correlations. Color of the labellum is correlated with: inflorescence appearance (ER), exposure of rachis (ER), inflorescence bract type (ER), cincinnus capacity (ER), floral density (ER), coiling of the floral tube (ER), the ratio of relative lengths of bract and calyx (ARD), the color of labellum blotch (ARD), and exsertion of stigma (ARD). Whereas, inflorescence bract type is correlated with: inflorescence appearance (ARD), cincinnus capacity (ER), number of flowers open per inflorescence per day (ER), coiling of the floral tube (ARD), type of labellum claw (ARD), the color of the labellum (ER), and exsertion of stigma (ARD). All the other correlations are highlighted in Appendix S1b and S1c.

Of the 13 categorical characters tested for phylogenetic signal using Fritz-Purvis’ D statistic, only three characters were found to have strong phylogenetic structure, four characters were found to have moderate ecological effects, and the remaining six characters did not reveal any effect (Appendix S1d). The D-statistic estimated for characters such as slender cylindrical inflorescence [E(D) = -0.90; Probability of E(D) resulting from Brownian phylogenetic structure > 0.95], cincinnus capacity [E(D) = -0.73; Probability of E(D) resulting from Brownian phylogenetic structure > 0.95], and coiling of floral tube [E(D) = -1.03; Probability of E(D) resulting from Brownian phylogenetic structure > 0.95] revealed a strong phylogenetic dependence or higher Brownian phylogenetic structure (Appendix S1d). In contrast, characters such as the number of flowers open per inflorescence per day [E(D) = 0.78; Probability of E(D) resulting from no (random) phylogenetic structure < 0.01], type of labellum claw [E(D) = 0.71; Probability of E(D) resulting from no (random) phylogenetic structure < 0.05], color of labellum blotch [E(D) = 0.93; Probability of E(D) resulting from no (random) phylogenetic structure < 0.01], and exsertion of stigma [E(D) = 0.80; Probability of E(D) resulting from no (random) phylogenetic structure < 0.05] revealed moderate ecological effects (Appendix S1d).

Of the 13 categorical characters tested for phylogenetic signal using δ-statistic, eight characters were found to have strong phylogenetic structure (Appendix S1d). The characters that showed higher phylogenetic signal were slender cylindrical inflorescence, rachis exposure, inflorescence bract type, floral density, cincinnus capacity, relative lengths of bract and calyx, coiling of floral tube and labellum colour.

### Phylogenetic signal and character correlation among continuous characters

The mapping of eight continuous characters revealed trait values (Appendix S2b) ranging between 1.7 (anther length; Appendix S2bH) and 15 (floral tube length; Appendix S2bF).

Both Pagel’s λ and Blomberg’s K were estimated for testing the hypothesis of phylogenetic signal variation among continuous floral characters in *Hedychium*. Three characters-lateral staminode length, floral tube length and anther length showed strong phylogenetic signal using the Pagel’s λ. The phylogenetic signals estimated using Blomberg’s K were weaker overall.

Of the eight continuously variable characters tested in the present study, five (labellum length, labellum width, lateral staminode length, floral tube length, anther length) are shown to possess significant phylogenetic signal according to Pagel’s λ (significantly different from 0; *P*-value < 0.01), and four (labellum length, labellum width, lateral staminode length, floral tube length) are shown to possess significant phylogenetic signal according to Blomberg’s K statistics (significantly different from 1; *P*-value < 0.01) based on absolute variables (Appendix S1e). Pagel’s λ ranged from 0.37–0.73; whilst Blomberg’s K ranged from 0.24–0.4. Notice that the Pagel’s λ values are significantly different from 1 in four continuous characters based on absolute variables (*P*-value < 0.01), indicating that although there is a considerable phylogenetic signal, other evolutionary processes might be playing a role rather than the pure Brownian motion in these traits. Fitting four different evolutionary models to the data further revealed that the assumption of no phylogenetic signal in traits under the WN model is rejected (OU ∼ WN: pchisq < 0.05; Appendix S1e; the OU model is favored over BM and better explains the evolutionary trend of labellum length, labellum width, labellum notch depth, lateral staminode length, floral tube length and anther length (OU ∼ BM: pchisq < 0.05). The rate of character evolution (Sig^2^) under the BM model ranged from 0.04 to 3.14, with the highest in floral tube length and lowest in anther length whereas the rate of character evolution (Sig^2^) under the OU model ranged from 0.06 to 16.79, with the highest in filament length and lowest in labellum notch depth (Appendix S1e). In the OU model, the alpha parameter varied from 0.27 in floral tube length to 0.88 in labellum width. It is already inferred that as the values of alpha parameters become larger, the effects of changing alpha on model predictions are increasingly smaller and large values of alpha are indistinguishable from the WN model (Cooper et al., 2016). Among all the continuous characters tested, lateral staminode width and filament length were the only two characters that favoured the White Noise model over the OU model, with alpha values greater than one.

PGLS analyses indicated a moderate correlation between seven traits and labellum width (Appendix S1e), with adjusted R^2^ ranging from 0.16 to 0.58. Overall, 81% of the variance found in labellum width can be explained by the combination of the other seven variables. Conversely, a relatively weak correlation was demonstrated between floral tube length and most individual traits, with R^2^ ranging from 0.03 (against labellum width) to 0.18 (against lateral staminode length) and the combination of the other five variables only reflecting 18% of lateral staminode length variance. Remarkably, there was weak correlation between labellum width and floral tube length (*P*-value > 0.05) and a negative correlation with floral tube length and filament length (Appendix S1f, S1g and S1h).

The disparity through time analysis showed morphological dissimilarity among all eight continuous traits on a multidimensional scale (Appendix S2c). The phylomorphospace analysis revealed the clustering of taxa that formed distinct clades (Appendix S2d).

### Trait dependent diversification in Hedychium

The hisse analysis to understand (categorical) trait-dependent diversification within the genus revealed that the floral density, cincinnus capacity, number of flowers open per inflorescence per day, color of the labellum and labellum blotch color were associated with the diversification of *Hedychium*. Whereas the inflorescence appearance, exposure of inflorescence rachis, bract type, bract to calyx length ratio, floral tube coiling were known to affect the diversification of the genus with a hidden state (Appendix S1i). Characters such as slender cylindrical inflorescence, type of labellum claw and exsertion of stigma revealed a character-independent fashion in the diversification of the genus. The best model favoured for each of the distinct discrete traits are summarised in Appendix S1i.

The BAMM analysis carried out to understand the phenotypic evolution among continuous traits revealed that both labellum width (Fig. 3) and floral tube length (Fig. 4) showed distinct diversification rate dynamics in the clade IV, the largest clade in the phylogeny of *Hedychium*. The other six traits did not reveal any significant difference in their diversification rate dynamics, in a clade-dependent fashion (Appendix S2e).

**FIGURE 3.**
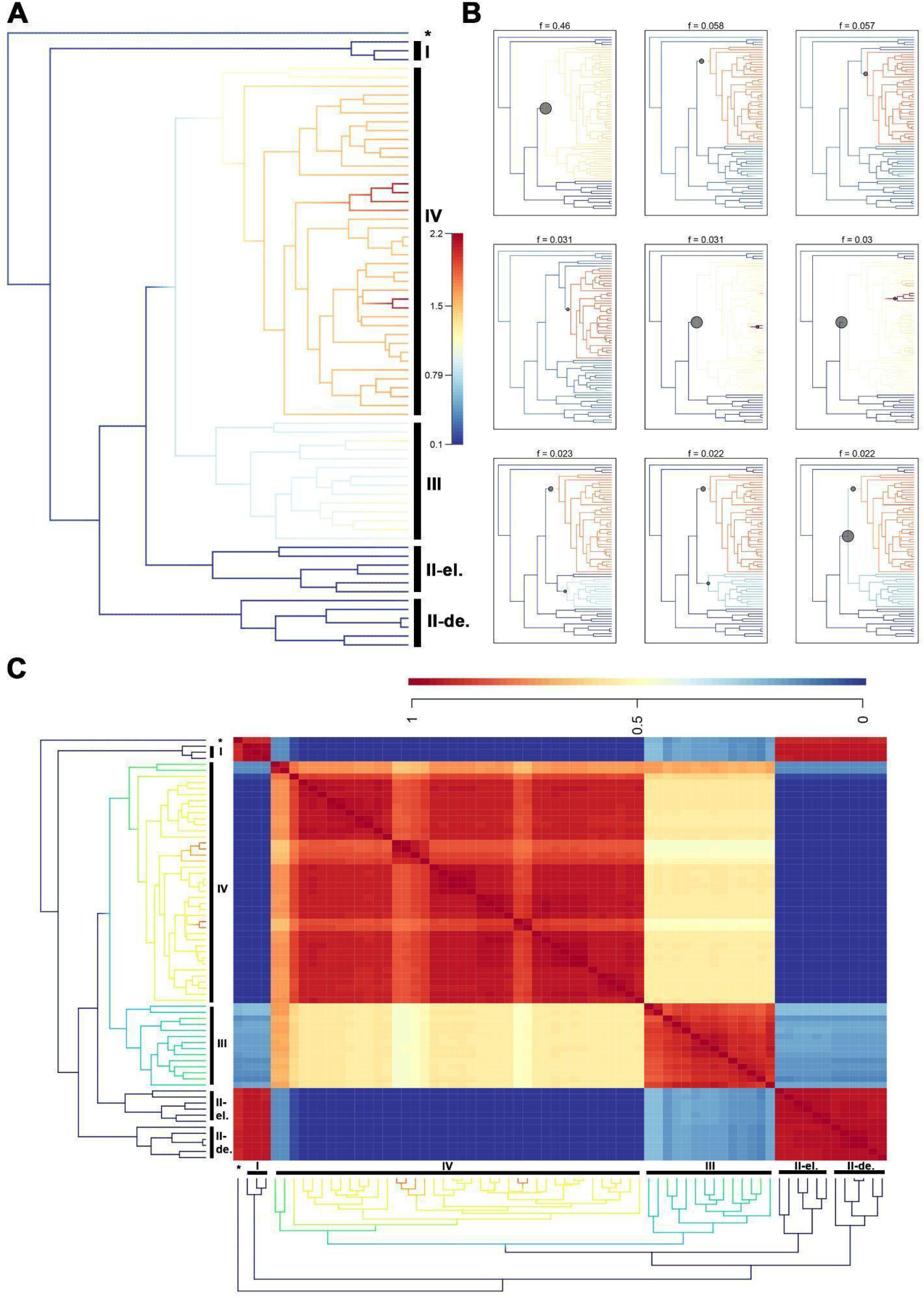
Phenotypic evolution of labellum width in *Hedychium*. (A) Mean phylorate plot. (B) Nine most probable macroevolutionary rate shift configurations (and their overall frequencies). (C) Macroevolutionary cohort matrix.

**FIGURE 4.**
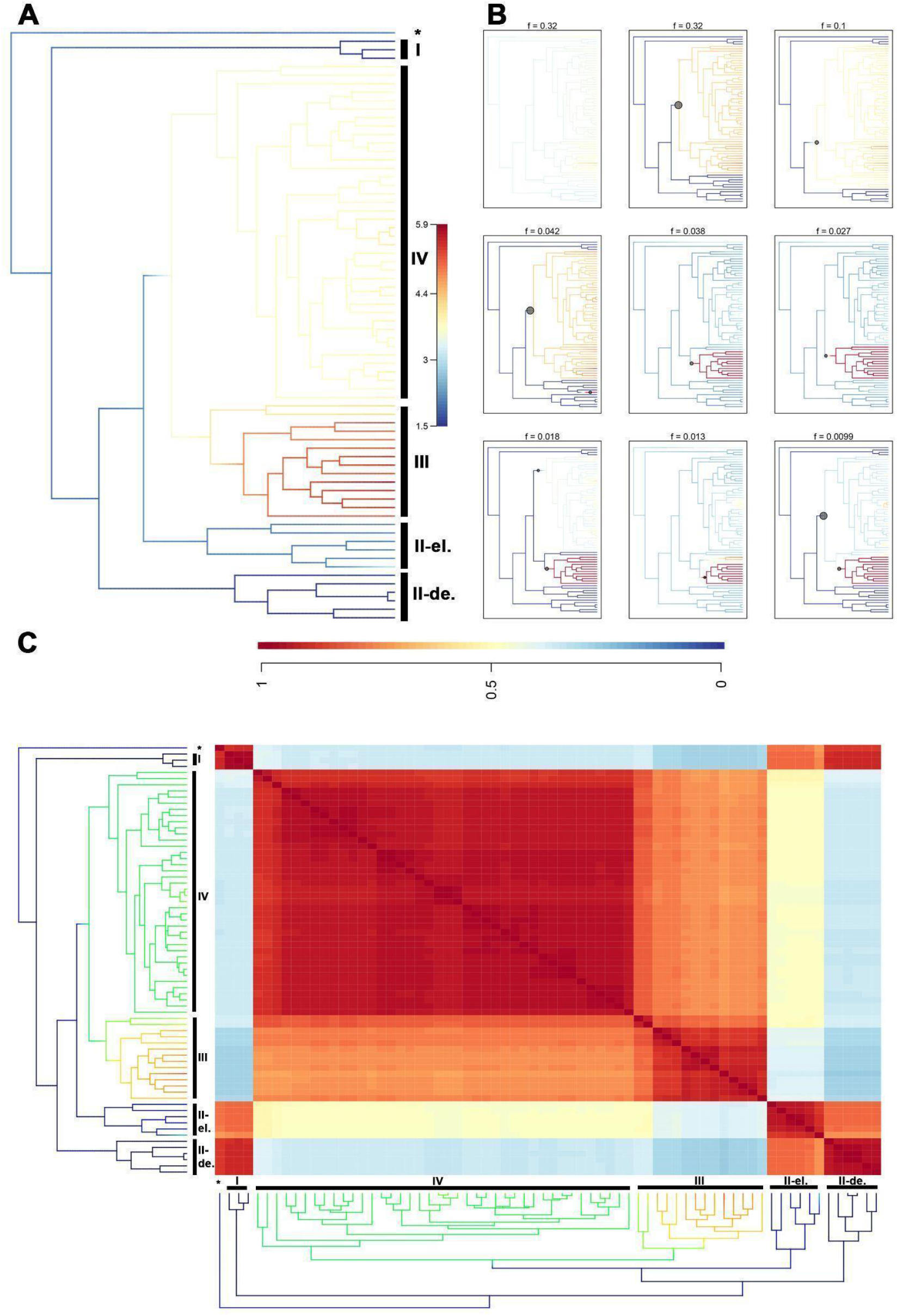
Phenotypic evolution of floral tube length in *Hedychium*. (A) Mean phylorate plot. (B) Nine most probable macroevolutionary rate shift configurations (and their overall frequencies). (C) Macroevolutionary cohort matrix.

## DISCUSSION

### Synapomorphies and clade-defining floral characters in Hedychium

Among the 13 categorical inflorescence and floral characters tested for putative synapomorphies in *Hedychium*, we identified five characters, namely, cincinnus capacity, inflorescence bract type, slender cylindrical inflorescence, coiling of floral tube, and ratio of bract and calyx lengths, to be phylogenetically informative.

### Cincinnus capacity and bract type

Both cincinnus capacity and inflorescence bract type were identified to be useful in distinguishing all or most of the clades of *Hedychium*. Cincinnus capacity defines the total number of flowers present in a bract and to that effect also represents the total flowers present in an inflorescence. Bract type refers to the folded or open nature of the inflorescence bracts, which bear flowers. The number of flowers per bract (cincinnus capacity as defined here) and the bract type have been mentioned in all major treatments of the genus (Wallich, 1853; Baker, 1892). Wood et al. (2000) was the first to attempt mapping cincinnus capacity onto a phylogenetic tree and they concluded that it was the only synapomorphic character for the genus. While Wood et al. (2000) did not use bract type to define the clades, it has been observed that bract type may be an important character to define clades (Sanoj, 2011), but this was never tested on a phylogenetic scale, until our study. Our results concur that clades in *Hedychium* can be defined by cincinnus capacity as well as bract type, with the exception being clade IV. Bract type was used by Schumann (1904) for classifying *Hedychium*, lending support to the utility of this character in taxonomic identifications. However, inflorescence bract type revealed a larger ambiguity within clade IV, which is both morphologically and taxonomically the most diverse clade, and phylogenetically the most derived clade (Ashokan et al., in review). Taxa included in clades I, II-el., and II-de. have only one or two flowers per bract whereas clades III and IV are made up of taxa with multi-flowered cincinni. Further, from the ancestral character-state reconstruction, we identified that clades I, II-el., II-de., and III, and the early diverging *H. pauciflorum* can be distinguished using inflorescence bract type and exposure of inflorescence rachis (the second character that was used by Schumann (1904) for classifying *Hedychium*). The folded bract type is found to be the ancestral form and is common among clades I, II-el., and II-de, while most taxa in clade III are characterised by open, imbricating bracts. We also found an intermediate bract type (partially folded bract) which was exclusive to *H. greenii* W.W.Sm. (clade IV) suggesting that this is an autapomorphic character.

### Features defining Clade III

Our previous studies have shown that taxa in clade III are predominantly epiphytic and island-dwelling (Ashokan et al., in review). Clade III taxa were also found to exhibit island dwarfism where the plant height is usually less than one meter with a reduction in the number of leaves in each shoot (Ashokan et al., in review). The clade III was supported by three synapomorphic characters: slender cylindrical inflorescence, coiling of floral tube, and shorter bract to calyx ratio (Fig. 1, Fig. 2). *Hedychium* is characterized by cylindrical inflorescence that show variation in their sizes, and flowers that have long floral tubes. Clade III is defined by slender cylindrical inflorescence which is characterized by the closely appressed nature of inflorescence bracts, with an apparent exsertion of floral tubes more than half its length. The exsertion of the floral tube often results in its coiling and notably we found these two characters to be correlated. From the phylogenetic signal estimation, both the slender cylindrical inflorescence and coiling of floral tubes were found to have strong phylogenetic structure although these characters do not define other clades within the genus.

### Floral trait evolution and pollination syndromes in Hedychium

#### Floral color

Floral color has been identified as one of the major cues in the description of pollination syndromes in *Hedychium* (Specht et al., 2012). Within *Hedychium* flowers, the labellum forms the most prominent structure and its color largely guides the taxonomic description of floral color in the genus, except in species where the labellum is reduced such as *H. horsfieldii* R.Br. ex Wall. and *H. tenellum* (K.Schum.) R.M.Sm. Primarily, the presence of white and pale flowers with fragrance has been mentioned as representing nocturnal traits (moth pollination, Darwin Correspondence Project, “Letter no. 9372”) and colored flowers without fragrance as diurnal traits (bird pollination, Specht et al., 2012). However these traits have never been used in any systematic treatment at the generic level (all studies from Schumann, 1904 to Wood et al., 2000). Our results show that flowers with bright colors (red and orange) have evolved multiple times within the genus, and most brightly colored flowers are concentrated in clade IV, with red and pink flowered taxa found only in this clade. Among the clades of *Hedychium* that show the presence of brightly colored flowers, we observed that non-white flowers evolved at least once (clades II-de. and III) or repeatedly as observed in clade IV. Clades II-de. and III show rare occurrences of taxa with bright orange-colored flowers (∼ ≤ 2 taxa), for example, *H. densiflorum* in clade II-de. and *H. longicornutum* Griff. ex Baker in clade III. Although the ancestral floral color form in *Hedychium* was observed to be equivocal in our analyses (Fig. 2C), we observed that clades I, II-el., and III are dominated by taxa with only white or pale-colored flowers. Despite the lack of strong phylogenetic signal supporting association of floral color in the genus, the dominance of white or pale flowers (or rare occurrence of brightly colored flowers) in two of the early diverging clades (clades I, and II-el.) and the late diverging clade III suggest that the white or pale flowers are ancestral traits while the presence of colored flowers is recent and may be influenced by ecological factors such as pollinators (Ashokan, 2020; Saryan, 2021).

#### Patterns in floral trait evolution

Of the 13 categorical and eight continuous characters investigated here, three categorical characters (floral density, number of flowers open per inflorescence per day and labellum color) and four continuous characters (labellum length, labellum width, floral tube length and filament length) were identified to have informative evolutionary patterns that revealed the role of ecological effects in the diversification of *Hedychium*. In our character correlation analyses, we observed that there is a weak negative correlation between floral tube length and filament length, suggesting presence of phenotypic redundancy in overall nectar and pollen presentations among flowers of *Hedychium* spp. The BAMM analysis shows that both labellum width and floral tube length have distinct diversification rate dynamics within the clade IV, thus indicating that these characters were probably associated with the diversification of lineages included in this clade (Fig. 3, Fig. 4). Notably, these two characters revealed moderately strong phylogenetic signals and also displayed larger variations among clade IV taxa (Appendix S1d). Overall, the taxa representing clade IV are found to have a wide range of floral size and color and this pattern of high variation in floral traits can be speculated to be indicative of adaptive radiation (see Lunau, 2004).

To understand the occurrence of pollination syndromes and their evolutionary patterns we examined floral characters to identify possible phylogenetic constraints. From our pollination studies of *Hedychium* we classified all species into one of the following functional groups: i) night opening flowers (mostly white, and heavily fragrant between dusk and dawn) and ii) day opening flowers (mostly brightly colored, mildly fragrant or not at all, open after dawn). The ancestral character-state reconstructions revealed that the night opening flowers have evolved repeatedly in the genus, across all clades (Fig. 2C). Additionally, the remarkable variation in the floral tube length (∼ 1 to 10 cm) across white-flowered taxa suggests multiple transitions to different types of pollinators or pollination syndromes (A. Ashokan, pers. obs.). The bright-colored flowers are not common in *Hedychium* and were found to have evolved in most clades (except clades I, and II-el.). This also signifies that diurnal pollination appeared as a derived trait in the genus, with majority of the bright-colored flowers present in the most derived clade IV. Although this finding partially supports the inferences on pollination syndromes in *Hedychium* by Specht et al. (2012) as well as the proposal of a floral type (bird flowers) in the genus (based on ovule protection) by Grant (1950), more field observations are warranted for confirming the bird pollination in *Hedychium*.

Supporting our findings on the presence of multiple pollination syndromes within the genus, field studies carried out by Saryan et al. (in prep, TrEE lab, IISER Bhopal; Ashokan and Gowda, 2017) concluded that taxa from clades I, II-el., II-de., and IV host a wide variety of floral visitors ranging from bees to birds and flies to beetles, with most of them having the potential to contribute to pollination or larceny (nectar and pollen). Further, most *Hedychium* flowers remained open for more than 24 hours with abundant floral rewards (both nectar and pollen), thus facilitating a higher visitor and/or pollinator diversity across these taxa (Saryan et al., in prep.). Given our phylogenetic and ecological observations, we conclude that pollinator diversification within the genus is common and specializations in plant-pollinator interactions are rare.

Darwin’s prediction of pollinators for *Hedychium* (*H. coronarium, H. gardnerianum* and *H. coccineum*) based on his observation of floral traits (Darwin Correspondence Project) remains to be an unresolved story despite our analyses because taxa within clade IV (circum-Himalayan clade) are highly variable in their floral traits (both floral size and color), have low phylogenetic resolution within the clade, and pollinators were not found in most taxa. While syndrome-like features were observed in many taxa, factors such as flowering phenology, pollination ecology, and current state of anthropogenic disturbances in most of our field sites seem to have overshadowed our documentation of legitimate pollinators in this group (P. Saryan et al., in prep.). Thus, presence of either moth-pollination or bird-pollination could not be confirmed through field experiments or phylogenetic reconstructions. On the whole, *Hedychium* exists as another challenging system from the IMR presenting complexities to advocates of the biological species concept and the idea of reproductive isolation.

## CONCLUSIONS

Flowers are one of the key innovations among angiosperms that are taxonomically critical, under ecological selection (especially, pollinator selection), and an important indicator of plant fitness (fruit set and seed set). In our present study, we discuss floral evolution patterns within the genus *Hedychium* especially related to inflorescence shape, bract type and cincinnus capacity. We also present results from our investigations of the role of floral traits such as floral color and floral size (via labellum) and its lack of association with pollinators or pollination syndromes (moth and bird pollination). We found weak correlations among these characters and found floral color to mostly define early diverging clades which show dominance of white or pale colored flowers. The circum-Himalayan clade IV which is the most derived and speciose clade was observed to be the most complex clade with high variability in most of its floral characters. We conclude that *Hedychium* is a young messy genus with poorly defined species boundaries resulting probably from non-specific pollen transfer and interspecific compatibility (Saryan, 2021) further contributing to the creation of tension zones and taxonomic confusions across the genus (Ashokan, 2020). Our conclusive results on phylogenetic relationship and floral trait evolution within the genus and our preliminary results on the relationship between floral traits and pollination traits are of considerable value for future integrated studies that can incorporate floral fragrance in understanding *Hedychium*-pollinator interactions within the IMR. Overall, the central message from our study is that evolutionary patterns of gingers (Zingiberaceae), especially taxa that are exclusive to IMR, need to be studied using an integrated approach of both ecological and evolutionary studies at both local and regional scale.

## ACKNOWLEDGMENTS

We thank the respective Indian state forest departments for field research permits-Arunachal Pradesh, Kerala, Manipur, Meghalaya, Mizoram, and Nagaland. We also thank the following herbaria and associated botanic gardens - ARUN, ASSAM, BHPL, BM, BO, BSA, BSHC, BSI, CAL, E, HLA, K, LINN, LIV, MH, QBG, SING, TBGT, US, and W for letting us access the specimens in their care and allowing tissue sampling from their herbarium or living collections. We express our sincere gratitude to Colton Collins (Plant Group Hawaii), Ernst Vitek (W), Marlina Ardiyani (BO), Mathew Dan (TBGT), Nripemo Odyuo (ASSAM), and Sinjumol Thomas (RHT) for sharing some of the molecular grade leaf tissues. We also thank the CIPRES Science Gateway for providing a platform to conduct all our phylogenetic analyses. Finally, we are thankful to Aleena Xavier, Prasanna N.S., Preeti Saryan, and Saket Shrotrifor their support in field expeditions, sequencing, and R studio. AA acknowledges funds from the Council of Scientific and Industrial Research for the Senior Research Fellowship, the Heliconia Society International for a Field Research Grant, and the Sibbald Trust at the Royal Botanic Garden Edinburgh for a grant to visit UK herbaria; VG received funds from Science and Engineering Research Board, Ministry of Human Resource Development (Ministry of Education), National Geographic Society, and Department of Biotechnology towards this study, and we thank IISER Bhopal for the infrastructure. Research of JL-Š is supported by the National Parks Board, Singapore. The RBGE is supported by the Scottish Government’s Rural and Environmental Science and Analytical Services Division.

## AUTHOR CONTRIBUTIONS

**Ajith Ashokan:** Conceptualization, Methodology, Resources, Tissue collection, Voucher processing, Software, Data curation, Investigation, Writing - original draft, Visualization. **Piyakaset Suksathan:** Tissue collection, Voucher processing, Writing - commenting on the drafts and approving the final manuscript. **Jana Leong-Škorničková:** Tissue collection, Voucher processing, Writing - commenting on the drafts and approving the final manuscript. **Mark Newman:** Access to material, herbarium study at BM, E, LINN, K, Writing and commenting on the drafts and approving the final manuscript. **W. John Kress:** Conceptualization, tissue collection, writing. **Vinita Gowda:** Conceptualization, Methodology, Resources, Writing - original draft, Project administration, Funding acquisition.

## APPENDIX S2 Supplemental Figures for

**Appendix S2a.**
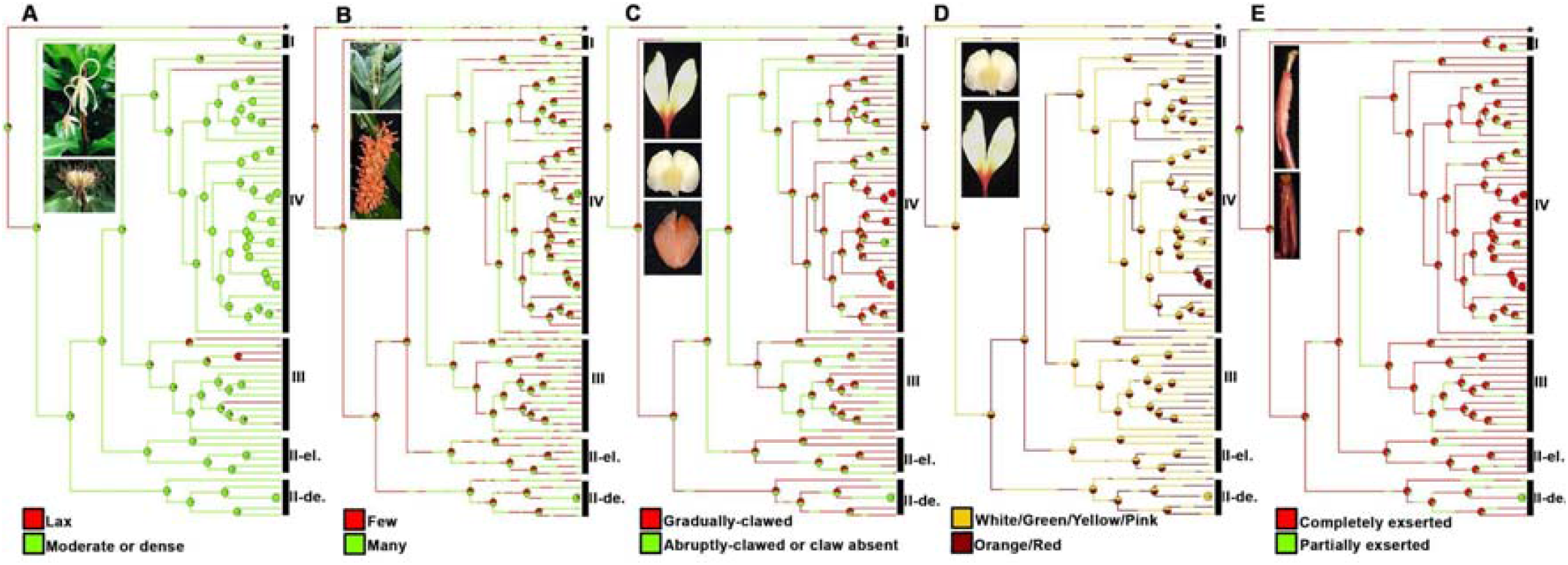
Ancestral character-state reconstructions of discrete floral traits in *Hedychium*. (A) Floral density, (B) Number of flowers open per inflorescence per day, (C) Type of claw, (D) color of labellum blotch, and (E) exsertion of stigma.

**Appendix S2b.**
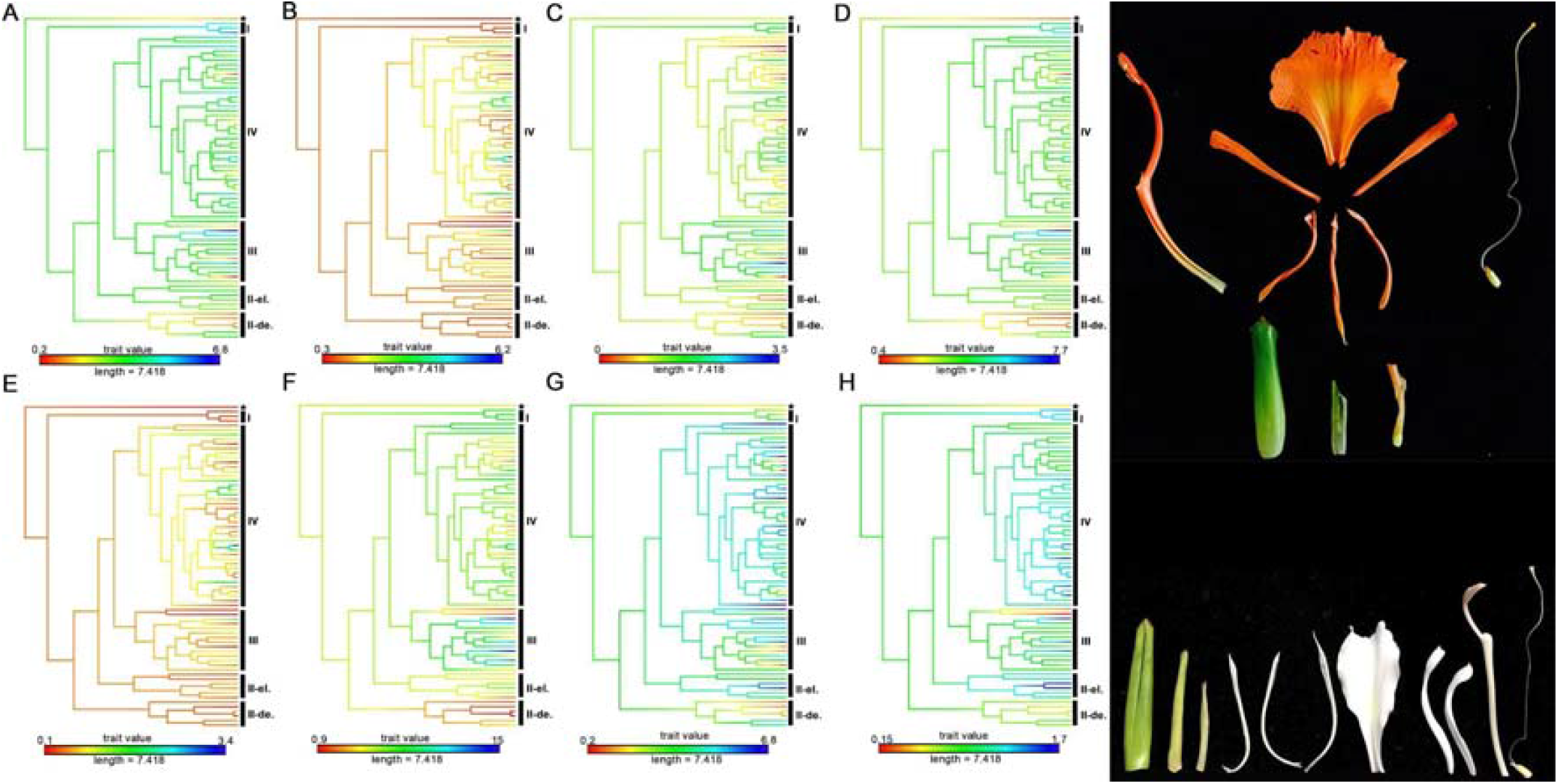
Continuous mapping of eight floral characters in *Hedychium*. (A) Labellum length, (B) Labellum width, (C) Labellum notch depth, (D) Lateral staminode length, (E) Lateral staminode width, (F) Floral tube length, (G) Filament length, and (H) Anther length.

**Appendix S2c.**
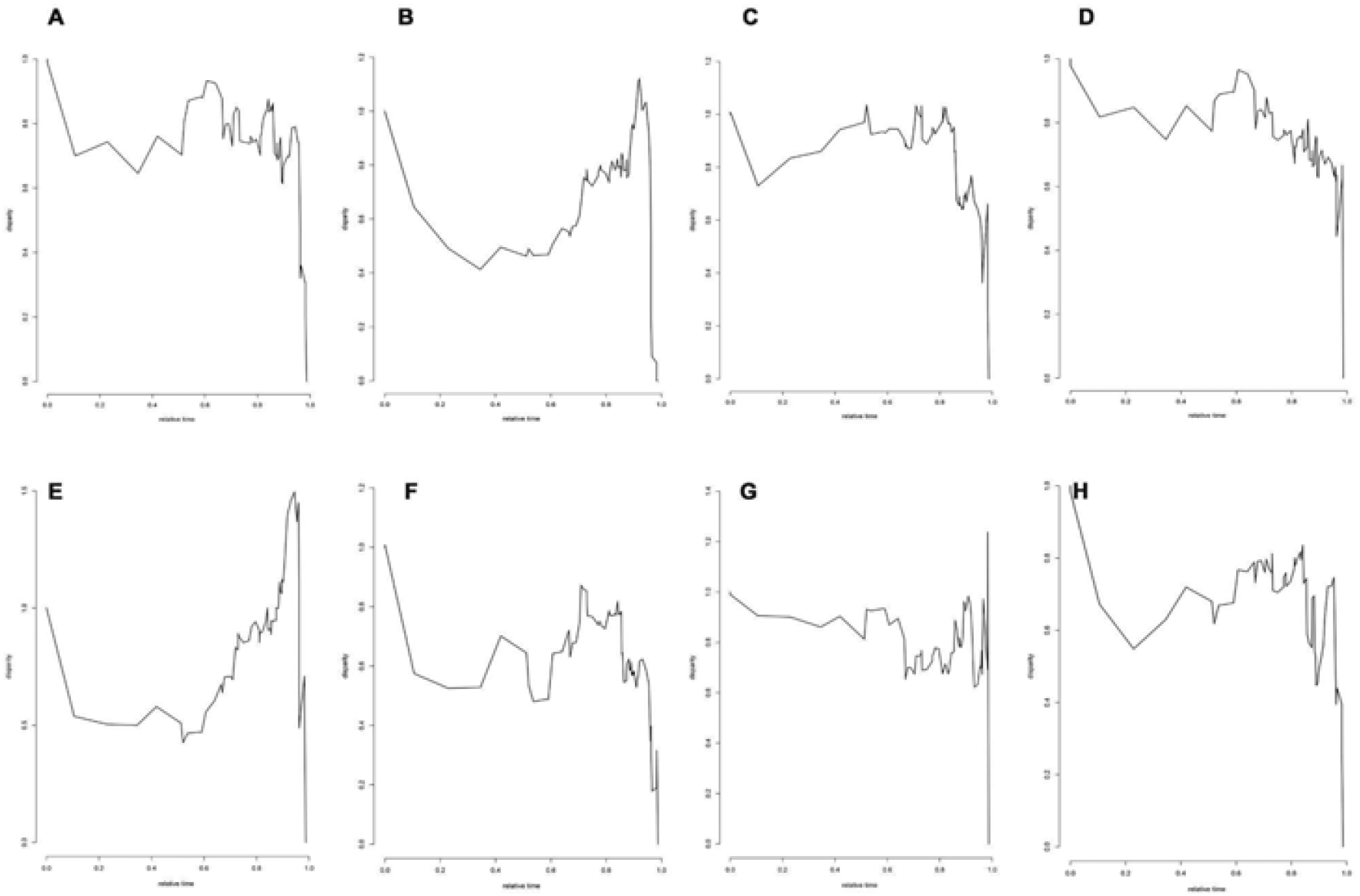
Disparity through time analysis for the eight floral characters in *Hedychium*. (A) Labellum length, (B) Labellum width, (C) Labellum notch depth, (D) Lateral staminode length, (E) Lateral staminode width, (F) Floral tube length, (G) Filament length, and (H) Anther length.

**Appendix S2d.**
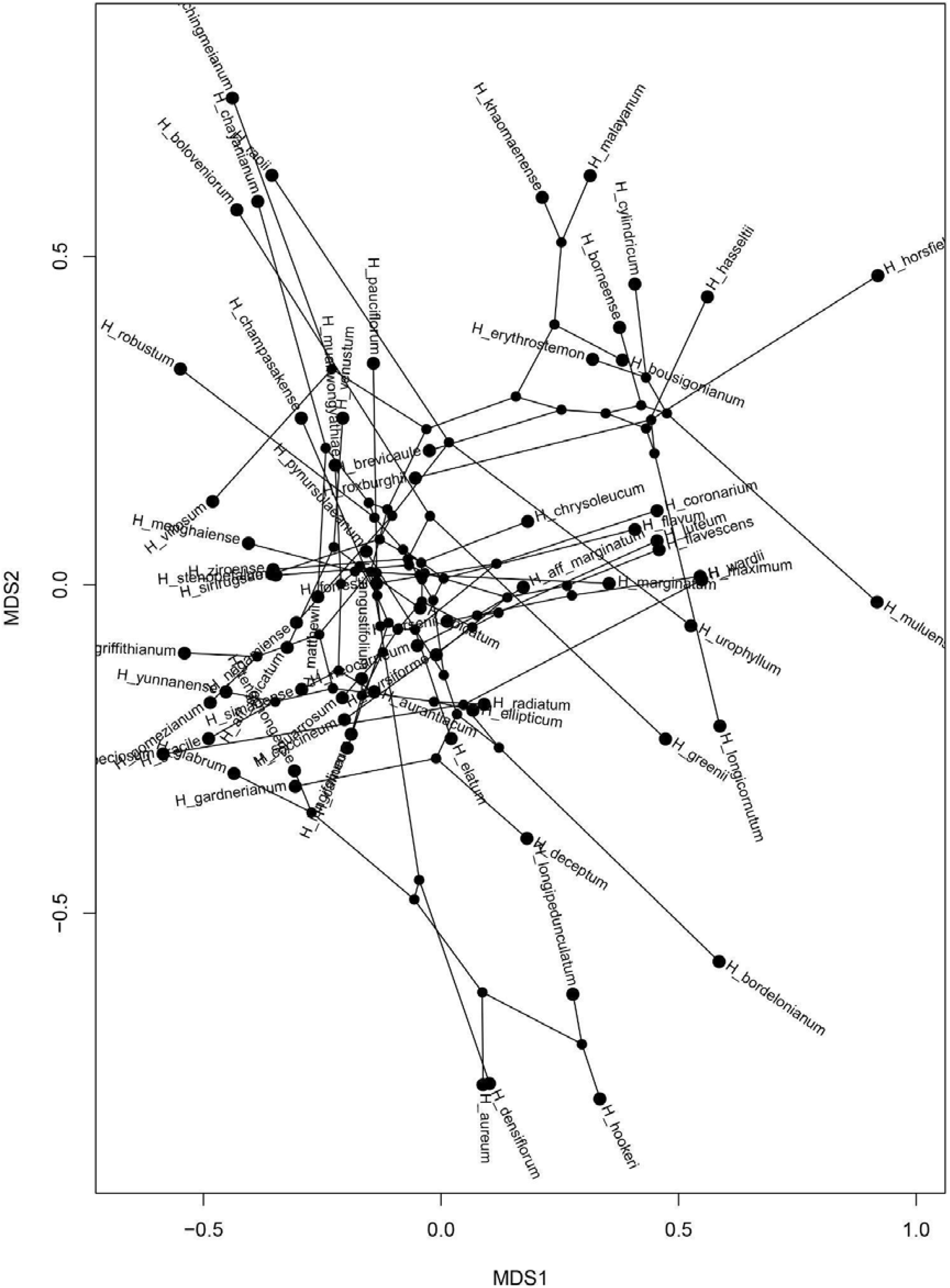
Phylomorphospace analysis for the thirteen discrete and eight continuous floral characters in *Hedychium*.

**Appendix S2e.**
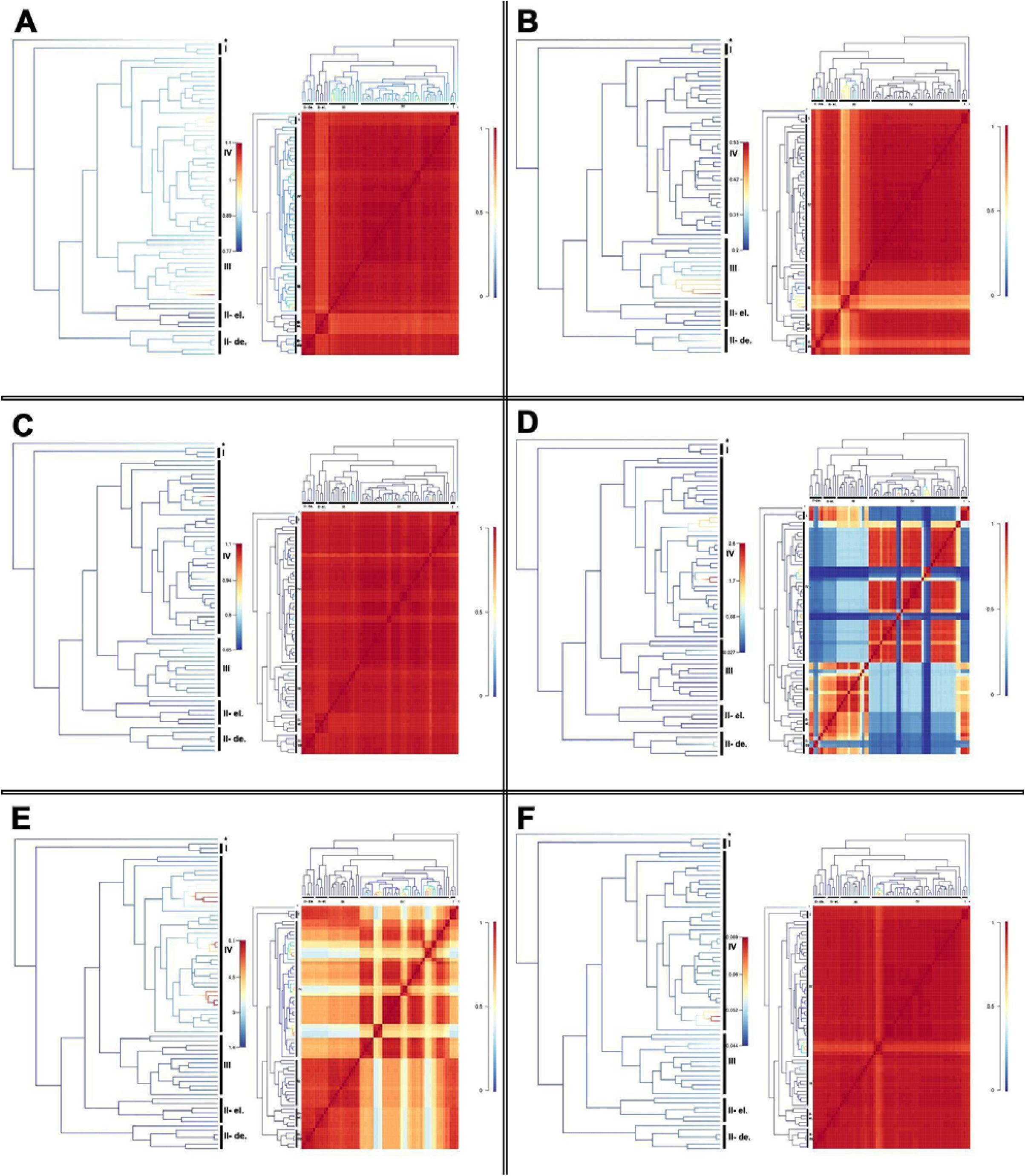
Phenotypic evolution of (A) Labellum length, (B) Labellum notch depth, (C) Lateral staminode length, (D) Lateral staminode width, (E) Filament length, and (G) Anther length in the diversification of *Hedychium*.

