## Appendix S1 for "Floral evolution and pollinator diversification in *Hedychium* J.Koenig (Zingiberaceae): one of Mr. Darwin’s tropical fantasies"

**APPENDIX S1 Supplemental Tables for**

**Floral evolution and pollinator diversification in *Hedychium* J.Koenig (Zingiberaceae): one of Mr. Darwin’s tropical fantasies**

Ajith Ashokan ^1, a^, Piyakaset Suksathan ^2^, Jana Leong-Škorničková ^3^, Mark Newman ^4^, W. John Kress ^5^, Vinita Gowda ^1,b^

^1^ Tropical Ecology and Evolution (TrEE) Lab, Department of Biological Sciences, Indian Institute of Science Education and Research (IISER) Bhopal, Madhya Pradesh 462066, India

^2^ Herbarium (QBG), Queen Sirikit Botanic Garden, P. O. Box 7, Mae Rim, Chiang Mai 50180, Thailand

^3^ Research & Conservation branch, Singapore Botanic Gardens, 1 Cluny Road, 259569, Singapore

^4^ Royal Botanic Garden Edinburgh, 20A Inverleith Row, Edinburgh EH3 5LR, Scotland, United Kingdom

^5^ Department of Botany, MRC-166, National Museum of Natural History, Smithsonian Institution, P. O. Box 37012, Washington, DC 20013-7012, United States

### **TABLE OF CONTENTS**

**Appendix	Page**

**Appendix S1a	3**

**Appendix S1b	5**

**Appendix S1c	18**

**Appendix S1d	20**

**Appendix S1e	21**

**Appendix S1f	24**

**Appendix S1g	28**

**Appendix S1h	29**

**Appendix S1i	30**

### **Appendix S1a.** Comparison of different models used in the ancestral character-state reconstruction of discrete binary characters in *Hedychium*.

| **Sl.no.** | **Code** | **Character** | **# States** | **Character states** | **LnL for ER** | **AIC for ER** | **AICc for ER** | **LnL for SYM** | **AIC for SYM** | **AICc for SYM** | **LnL for ARD** | **AIC for ARD** | **AICc for ARD** |
| --- | --- | --- | --- | --- | --- | --- | --- | --- | --- | --- | --- | --- | --- |
| 1 | slender | Slender cylindrical inflorescence | 2 | Absent (0), Present (1) | -16.22024 | 34.44048 | **34.49928** | -16.22024 | 38.44048 | 38.80408 | -12.62229 | 37.24458 | 38.57788 |
| 2 | appear | Inflorescence appears wider than long | 2 | No (0), Yes (1) | -39.03065 | 80.0613 | **80.1201** | -39.03065 | 84.0613 | 84.4249 | -34.06973 | 80.13946 | 81.47276 |
| 3 | rachis | Rachis exposure | 2 | Fully or partially exposed (0), Not exposed (1) | -44.19496 | 90.38992 | **90.44872** | -44.19496 | 94.38992 | 94.75352 | -43.86326 | 99.72652 | 101.0598 |
| 4 | bract | Inflorescence bract type | 2 | Folded or partially folded (0), Open (1) | -38.33147 | 78.66294 | **78.72174** | -38.33147 | 82.66294 | 83.02654 | -37.28557 | 86.57114 | 87.90444 |
| 5 | density | Floral density | 2 | Lax (0), Moderately or highly dense (1) | -33.40199 | 68.80398 | **68.86278** | -33.40199 | 72.80398 | 73.16758 | -32.28612 | 76.57224 | 77.90554 |
| 6 | cincinnus | Cincinnus capacity | 2 | One or two (0), More than two (1) | -22.93766 | 47.87532 | **47.93412** | -22.93766 | 51.87532 | 52.23892 | -22.92614 | 57.85228 | 59.18558 |
| 7 | flowers | Flowers open per day | 2 | Few (0), Many or the whole inflorescence open simultaneously (1) | -47.6425 | 97.285 | **97.3438** | -47.6425 | 101.285 | 101.6486 | -47.56589 | 107.1318 | 108.4651 |
| 8 | calyx | Relative length of bract and calyx | 2 | More or equal (0), Less (1) | -34.31604 | 70.63208 | **70.69088** | -34.31604 | 74.63208 | 74.99568 | -33.36997 | 78.73994 | 80.07324 |
| 9 | coiling | Floral tube coiling | 2 | No (0), Yes (1) | -14.60833 | 31.216 | **31.2748** | -14.60833 | 35.2166 | 35.5802 | -14.34394 | 40.6878 | 42.0211 |
| 10 | claw | Labellum claw | 2 | Gradually clawed (0), Absent or Abruptly clawed (1) | -46.50977 | 95.01954 | **95.07834** | -46.50977 | 99.01954 | 99.38314 | -42.60072 | 97.20144 | 98.53474 |
| 11 | labellum | Labellum color | 4 | White (0), Yellow (1), Orange (2), Red or Pink (3) | -84.28136 | 170.5627 | 170.6215 | -76.85573 | 159.7115 | 160.0751 | -66.71308 | 145.4262 | **146.7595** |
| 12 | blotch | Labellum blotch color | 2 | White/Green/Yellow/Pink (0), Orange or Red (1) | -47.62911 | 97.25822 | **97.31702** | -47.62911 | 101.2582 | 101.6218 | -47.43456 | 106.8691 | 108.2024 |
| 13 | stigma | Stigma position | 2 | Completely exserted (0), Partially exserted (1) | -43.18099 | 88.36198 | 88.42078 | -43.18099 | 92.36198 | 92.72558 | -37.28874 | 86.57748 | **87.91078** |

### **Appendix S1b.** Comparison of different models used in the character correlation analyses of discrete binary characters in *Hedychium*.

| **Sl. no.** | **Model** | **Character combinations** | **Independent model (LnL)** | **Independent model (AIC)** | **Dependent model (LnL)** | **Dependent model (AIC)** | **Likelihood ratio** | **p-value** |
| --- | --- | --- | --- | --- | --- | --- | --- | --- |
| 1a | ER | bract vs density | -72.42954 | 148.8591 | -70.87093 | 149.7419 | 3.11722 | 0.210428 |
| 1b | SYM | bract vs density | -72.42954 | 148.8591 | -70.87093 | 149.7419 | 3.11722 | 0.210428 |
| 1c | ARD | bract vs density | -70.93291 | 149.8658 | -68.57364 | 153.1473 | 4.71854 | 0.317415 |
| **2a** | **ER** | **bract vs cincinnus** | **-61.61658** | **127.2332** | **-58.33843** | **124.6769** | **6.55631** | **0.0376978** |
| 2b | SYM | bract vs cincinnus | -61.61658 | 127.2332 | -58.33843 | 124.6769 | 6.55631 | 0.0376978 |
| 2c | ARD | bract vs cincinnus | -60.96967 | 129.9393 | -56.66709 | 129.3342 | 8.60517 | 0.0717628 |
| **3a** | **ER** | **bract vs flowers** | **-86.93471** | **177.8694** | **-79.97385** | **167.9477** | **13.9217** | **0.000948282** |
| 3b | SYM | bract vs flowers | -86.93471 | 177.8694 | -79.97385 | 167.9477 | 13.9217 | 0.000948282 |
| 3c | ARD | bract vs flowers | -86.21268 | 180.4254 | -76.70106 | 169.4021 | 19.0232 | 0.000777725 |
| 4a | ER | bract vs claw | -85.80198 | 175.6040 | -81.61587 | 171.2317 | 8.37223 | 0.0152053 |
| 4b | SYM | bract vs claw | -85.80198 | 175.6040 | -81.61587 | 171.2317 | 8.37223 | 0.0152053 |
| **4c** | **ARD** | **bract vs claw** | **-81.24751** | **170.4950** | **-74.46594** | **164.9319** | **13.5632** | **0.00882808** |
| **5a** | **ER** | **bract vs labellum*** | **-85.93739** | **175.8748** | **-82.74779** | **173.4956** | **6.37921** | **0.0411881** |
| 5b | SYM | bract vs labellum | -85.93739 | 175.8748 | -82.74779 | 173.4956 | 6.37921 | 0.0411881 |
| 5c | ARD | bract vs labellum | -81.00632 | 170.0126 | -78.06973 | 172.1395 | 5.87316 | 0.208823 |
| 6a | ER | bract vs blotch | -86.92132 | 177.8426 | -86.83254 | 181.6651 | 0.177563 | 0.915045 |
| 6b | SYM | bract vs blotch | -86.92132 | 177.8426 | -86.83254 | 181.6651 | 0.177563 | 0.915045 |
| 6c | ARD | bract vs blotch | -86.08136 | 180.1627 | -85.07276 | 186.1455 | 2.0172 | 0.732595 |
| 7a | ER | bract vs stigma | -82.34477 | 168.6895 | -74.14820 | 156.2964 | 16.3931 | 0.000275597 |
| 7b | SYM | bract vs stigma | -82.34477 | 168.6895 | -74.14820 | 156.2964 | 16.3931 | 0.000275597 |
| **7c** | **ARD** | **bract vs stigma** | **-75.93553** | **159.8711** | **-69.48018** | **154.9604** | **12.9107** | **0.0117205** |
| 8a | ER | density vs cincinnus | -56.84800 | 117.6960 | -55.46041 | 118.9208 | 2.77518 | 0.249676 |
| 8b | SYM | density vs cincinnus | -56.84800 | 117.6960 | -55.46041 | 118.9208 | 2.77518 | 0.249676 |
| 8c | ARD | density vs cincinnus | -55.99528 | 119.9906 | -53.64556 | 123.2911 | 4.69943 | 0.31955 |
| 9a | ER | density vs flowers | -82.16612 | 168.3322 | -79.85777 | 167.7155 | 4.61671 | 0.0994249 |
| 9b | SYM | density vs flowers | -82.16612 | 168.3322 | -79.85777 | 167.7155 | 4.61671 | 0.0994249 |
| 9c | ARD | density vs flowers | -81.23829 | 170.4766 | -78.05044 | 172.1009 | 6.37572 | 0.172792 |
| 10a | ER | density vs claw | -81.03339 | 166.0668 | -78.66410 | 165.3282 | 4.7386 | 0.0935464 |
| 10b | SYM | density vs claw | -81.03339 | 166.0668 | -78.66410 | 165.3282 | 4.7386 | 0.0935464 |
| 10c | ARD | density vs claw | -76.27313 | 160.5463 | -74.66759 | 165.3352 | 3.21107 | 0.523145 |
| **11a** | **ER** | **density vs labellum** | **-81.16881** | **166.3376** | **-74.97435** | **157.9487** | **12.3889** | **0.00204071** |
| 11b | SYM | density vs labellum | -81.16881 | 166.3376 | -74.97435 | 157.9487 | 12.3889 | 0.00204071 |
| 11c | ARD | density vs labellum | -76.03193 | 160.0639 | -71.54812 | 159.0962 | 8.96762 | 0.0619139 |
| 12a | ER | density vs blotch | -82.15274 | 168.3055 | -80.70076 | 169.4015 | 2.90396 | 0.234106 |
| 12b | SYM | density vs blotch | -82.15274 | 168.3055 | -80.70076 | 169.4015 | 2.90396 | 0.234106 |
| 12c | ARD | density vs blotch | -81.10697 | 170.2139 | -77.84955 | 171.6991 | 6.51485 | 0.163857 |
| **13a** | **ER** | **density vs stigma** | **-77.57618** | **159.1524** | **-73.33039** | **154.6608** | **8.4916** | **0.0143243** |
| 13b | SYM | density vs stigma | -77.57618 | 159.1524 | -73.33039 | 154.6608 | 8.4916 | 0.0143243 |
| 13c | ARD | density vs stigma | -70.96114 | 149.9223 | -67.71946 | 151.4389 | 6.48337 | 0.165841 |
| 14a | ER | cincinnus vs flowers | -71.35317 | 146.7063 | -70.57182 | 149.1436 | 1.56269 | 0.45779 |
| 14b | SYM | cincinnus vs flowers | -71.35317 | 146.7063 | -70.57182 | 149.1436 | 1.56269 | 0.45779 |
| 14c | ARD | cincinnus vs flowers | -71.27505 | 150.5501 | -67.25936 | 150.5187 | 8.0314 | 0.0904348 |
| 15a | ER | cincinnus vs claw | -70.22044 | 144.4409 | -67.84477 | 143.6895 | 4.75133 | 0.0929525 |
| 15b | SYM | cincinnus vs claw | -70.22044 | 144.4409 | -67.84477 | 143.6895 | 4.75133 | 0.0929525 |
| 15c | ARD | cincinnus vs claw | -66.30989 | 140.6198 | -62.96990 | 141.9398 | 6.67996 | 0.153799 |
| **16a** | **ER** | **cincinnus vs labellum** | **-70.35585** | **144.7117** | **-64.26001** | **136.5200** | **12.1917** | **0.00225221** |
| 16b | SYM | cincinnus vs labellum | -70.35585 | 144.7117 | -64.26001 | 136.5200 | 12.1917 | 0.00225221 |
| 16c | ARD | cincinnus vs labellum | -66.06869 | 140.1374 | -60.48670 | 136.9734 | 11.164 | 0.0247817 |
| **17a** | **ER** | **cincinnus vs blotch** | **-71.33978** | **146.6796** | **-67.80724** | **143.6145** | **7.06508** | **0.0292306** |
| 17b | SYM | cincinnus vs blotch | -71.33978 | 146.6796 | -67.80724 | 143.6145 | 7.06508 | 0.0292306 |
| 17c | ARD | cincinnus vs blotch | -71.14373 | 150.2875 | -66.73248 | 149.4650 | 8.8225 | 0.0656925 |
| 18a | ER | cincinnus vs stigma | -66.76323 | 137.5265 | -65.05770 | 138.1154 | 3.41106 | 0.181676 |
| 18b | SYM | cincinnus vs stigma | -66.76323 | 137.5265 | -65.05770 | 138.1154 | 3.41106 | 0.181676 |
| 18c | ARD | cincinnus vs stigma | -60.99790 | 129.9958 | -57.54597 | 131.0919 | 6.90386 | 0.141057 |
| **19a** | **ER** | **flowers vs claw** | **-95.53856** | **195.0771** | **-87.72362** | **183.4472** | **15.6299** | **0.000403659** |
| 19b | SYM | flowers vs claw | -95.53856 | 195.0771 | -87.72362 | 183.4472 | 15.6299 | 0.000403659 |
| 19c | ARD | flowers vs claw | -91.55290 | 191.1058 | -85.09737 | 186.1947 | 12.9111 | 0.0117186 |
| 20a | ER | flowers vs labellum | -95.67397 | 195.3480 | -94.01753 | 196.0351 | 3.3129 | 0.190816 |
| 20b | SYM | flowers vs labellum | -95.67397 | 195.3480 | -94.01753 | 196.0351 | 3.3129 | 0.190816 |
| 20c | ARD | flowers vs labellum | -91.31170 | 190.6234 | -89.11131 | 194.2226 | 4.40078 | 0.354474 |
| 21a | ER | flowers vs blotch | -96.65791 | 197.3158 | -96.41315 | 200.8263 | 0.489505 | 0.782898 |
| 21b | SYM | flowers vs blotch | -96.65791 | 197.3158 | -96.41315 | 200.8263 | 0.489505 | 0.782898 |
| 21c | ARD | flowers vs blotch | -96.38674 | 200.7735 | -96.00044 | 208.0009 | 0.772601 | 0.942083 |
| 22a | ER | flowers vs stigma | -92.08135 | 188.1627 | -82.34494 | 172.6899 | 19.4728 | 5.90922e-05 |
| 22b | SYM | flowers vs stigma | -92.08135 | 188.1627 | -82.34494 | 172.6899 | 19.4728 | 5.90922e-05 |
| **22c** | **ARD** | **flowers vs stigma** | **-86.24092** | **180.4818** | **-77.93664** | **171.8733** | **16.6085** | **0.00230241** |
| 23a | ER | claw vs labellum | -94.54124 | 193.0825 | -91.64945 | 191.2989 | 5.78358 | 0.0554768 |
| 23b | SYM | claw vs labellum | -94.54124 | 193.0825 | -91.64945 | 191.2989 | 5.78358 | 0.0554768 |
| 23c | ARD | claw vs labellum | -86.34653 | 180.6931 | -83.17439 | 182.3488 | 6.3443 | 0.174869 |
| 24a | ER | claw vs blotch | -95.52518 | 195.0504 | -93.09072 | 194.1814 | 4.8689 | 0.0876458 |
| 24b | SYM | claw vs blotch | -95.52518 | 195.0504 | -93.09072 | 194.1814 | 4.8689 | 0.0876458 |
| 24c | ARD | claw vs blotch | -91.42158 | 190.8432 | -89.40144 | 194.8029 | 4.04026 | 0.400584 |
| 25a | ER | claw vs stigma | -90.94862 | 185.8972 | -72.30142 | 152.6028 | 37.2944 | 7.97301e-09 |
| 25b | SYM | claw vs stigma | -90.94862 | 185.8972 | -72.30142 | 152.6028 | 37.2944 | 7.97301e-09 |
| **25c** | **ARD** | **claw vs stigma** | **-81.27575** | **170.5515** | **-62.01950** | **140.0390** | **38.5125** | **8.78366e-08** |
| 26a | ER | labellum vs blotch | -95.66059 | 195.3212 | -93.35281 | 194.7056 | 4.61557 | 0.0994814 |
| 26b | SYM | labellum vs blotch | -95.66059 | 195.3212 | -93.35281 | 194.7056 | 4.61557 | 0.0994814 |
| **26c** | **ARD** | **labellum vs blotch** | **-91.18038** | **190.3608** | **-85.51253** | **187.0251** | **11.3357** | **0.0230393** |
| 27a | ER | labellum vs stigma | -91.08404 | 186.1681 | -89.85589 | 187.7118 | 2.45629 | 0.292835 |
| 27b | SYM | labellum vs stigma | -91.08404 | 186.1681 | -89.85589 | 187.7118 | 2.45629 | 0.292835 |
| **27c** | **ARD** | **labellum vs stigma** | **-81.03455** | **170.0691** | **-76.21328** | **168.4266** | **9.64253** | **0.0468994** |
| 28a | ER | blotch vs stigma | -92.06797 | 188.1359 | -91.04860 | 190.0972 | 2.03874 | 0.360822 |
| 28b | SYM | blotch vs stigma | -92.06797 | 188.1359 | -91.04860 | 190.0972 | 2.03874 | 0.360822 |
| 28c | ARD | blotch vs stigma | -86.10959 | 180.2192 | -83.74330 | 183.4866 | 4.73258 | 0.315853 |
| 29a | ER | slender vs appear | -45.72561 | 95.45123 | -45.72561 | 99.45123 | 7.83785e-10 | 1 |
| 29b | SYM | slender vs appear | -45.72561 | 95.45123 | -45.72561 | 99.45123 | 7.83785e-10 | 1 |
| 29c | ARD | slender vs appear | -45.72561 | 99.45123 | -45.72561 | 107.45123 | -7.83785e-10 | 1 |
| 30a | ER | slender vs rachis | -49.50407 | 103.0081 | -49.50407 | 107.0081 | 0 | 1 |
| 30b | SYM | slender vs rachis | -49.50407 | 103.0081 | -49.50407 | 107.0081 | 0 | 1 |
| 30c | ARD | slender vs rachis | -49.50407 | 107.0081 | -49.50407 | 115.0081 | 0 | 1 |
| 31a | ER | slender vs bract | -40.02629 | 84.05257 | -40.02629 | 88.05257 | 0 | 1 |
| 31b | SYM | slender vs bract | -40.02629 | 84.05257 | -40.02629 | 88.05257 | 0 | 1 |
| 31c | ARD | slender vs bract | -40.02629 | 88.05257 | -40.02629 | 96.05257 | -7.76893e-10 | 1 |
| 32a | ER | slender vs density | -50.06817 | 104.1363 | -48.25653 | 104.5131 | 3.62328 | 0.163386 |
| 32b | SYM | slender vs density | -50.06817 | 104.1363 | -48.25653 | 104.5131 | 3.62328 | 0.163386 |
| 32c | ARD | slender vs density | -46.14919 | 100.2984 | -42.74178 | 101.4836 | 6.8148 | 0.146005 |
| 33a | ER | slender vs cincinnus | -39.25521 | 82.51043 | -37.29558 | 82.59117 | 3.91926 | 0.14091 |
| 33b | SYM | slender vs cincinnus | -39.25521 | 82.51043 | -37.29558 | 82.59117 | 3.91926 | 0.14091 |
| 33c | ARD | slender vs cincinnus | -36.02993 | 80.05986 | -33.66723 | 83.33445 | 4.72541 | 0.31665 |
| 34a | ER | slender vs flowers | -53.74698 | 111.494 | -53.74698 | 115.494 | 0 | 1 |
| 34b | SYM | slender vs flowers | -53.74698 | 111.494 | -53.74698 | 115.494 | 0 | 1 |
| 34c | ARD | slender vs flowers | -53.74698 | 115.494 | -53.74698 | 123.494 | 2.96945e-06 | 1 |
| 35a | ER | slender vs claw | -63.44061 | 130.8812 | -62.36075 | 132.7215 | 2.15971 | 0.339644 |
| 35b | SYM | slender vs claw | -63.44061 | 130.8812 | -62.36075 | 132.7215 | 2.15971 | 0.339644 |
| 35c | ARD | slender vs claw | -56.30777 | 120.6155 | -54.93017 | 125.8603 | 2.75522 | 0.599588 |
| 36a | ER | slender vs labellum | -53.33349 | 110.667 | -53.33349 | 114.667 | 1.01039e-11 | 1 |
| 36b | SYM | slender vs labellum | -53.33349 | 110.667 | -53.33349 | 114.667 | 1.01039e-11 | 1 |
| 36c | ARD | slender vs labellum | -53.33349 | 114.667 | -53.33349 | 122.667 | -4.07283e-11 | 1 |
| 37a | ER | slender vs blotch | -64.55995 | 133.1199 | -63.71239 | 135.4248 | 1.69512 | 0.428459 |
| 37b | SYM | slender vs blotch | -64.55995 | 133.1199 | -63.71239 | 135.4248 | 1.69512 | 0.428459 |
| 37c | ARD | slender vs blotch | -61.14162 | 130.2832 | -60.38949 | 136.7790 | 1.50425 | 0.825888 |
| 38a | ER | slender vs stigma | -59.98340 | 123.9668 | -57.92882 | 123.8576 | 4.10915 | 0.128147 |
| 38b | SYM | slender vs stigma | -59.98340 | 123.9668 | -57.92882 | 123.8576 | 4.10915 | 0.128147 |
| 38c | ARD | slender vs stigma | -50.99579 | 109.9916 | -49.19823 | 114.3965 | 3.59512 | 0.463564 |
| 39a | ER | slender vs coiling | -19.70329 | 43.40658 | -19.56238 | 47.12476 | 0.281823 | 0.868566 |
| 39b | SYM | slender vs coiling | -19.70329 | 43.40658 | -19.56238 | 47.12476 | 0.281823 | 0.868566 |
| 39c | ARD | slender vs coiling | -19.70329 | 47.40658 | -19.70329 | 55.40658 | 0 | 1 |
| **40a** | **ER** | **slender vs calyx** | **-51.04244** | **106.08488** | **-45.37538** | **98.75076** | **11.3341** | **0.00345803** |
| 40b | SYM | slender vs calyx | -51.04244 | 106.08488 | -45.37538 | 98.75076 | 11.3341 | 0.00345803 |
| 40c | ARD | slender vs calyx | -47.06905 | 102.1381 | -44.66542 | 105.3309 | 4.80725 | 0.307652 |
| 41a | ER | appear vs rachis | -84.08439 | 172.1688 | -72.20087 | 152.4017 | 23.767 | 6.90328e-06 |
| 41b | SYM | appear vs rachis | -84.08439 | 172.1688 | -72.20087 | 152.4017 | 23.767 | 6.90328e-06 |
| **41c** | **ARD** | **appear vs rachis** | **-79.31838** | **166.6368** | **-64.66942** | **145.3388** | **29.2979** | **6.80019e-06** |
| 42a | ER | appear vs bract | -77.81123 | 159.6225 | -68.39761 | 144.7952 | 18.8272 | 8.16051e-05 |
| 42b | SYM | appear vs bract | -77.81123 | 159.6225 | -68.39761 | 144.7952 | 18.8272 | 8.16051e-05 |
| **42c** | **ARD** | **appear vs bract** | **-72.71652** | **153.4330** | **-61.61323** | **139.2265** | **22.2066** | **0.000182308** |
| 43a | ER | appear vs density | -64.28313 | 132.5663 | -64.28313 | 136.5663 | 6.15777e-08 | 1 |
| 43b | SYM | appear vs density | -64.28313 | 132.5663 | -64.28313 | 136.5663 | 6.15777e-08 | 1 |
| 43c | ARD | appear vs density | -64.28313 | 136.5663 | -64.28313 | 144.5663 | -6.15777e-08 | 1 |
| 44a | ER | appear vs cincinnus | -62.22971 | 128.4594 | -61.39549 | 130.7910 | 1.66843 | 0.434215 |
| 44b | SYM | appear vs cincinnus | -62.22971 | 128.4594 | -61.39549 | 130.7910 | 1.66843 | 0.434215 |
| 44c | ARD | appear vs cincinnus | -57.77889 | 123.5578 | -56.62942 | 129.2588 | 2.29895 | 0.68096 |
| 45a | ER | appear vs flowers | -87.54783 | 179.0957 | -84.97248 | 177.9450 | 5.15069 | 0.0761274 |
| 45b | SYM | appear vs flowers | -87.54783 | 179.0957 | -84.97248 | 177.9450 | 5.15069 | 0.0761274 |
| 45c | ARD | appear vs flowers | -83.02191 | 174.0438 | -79.60697 | 175.2139 | 6.82986 | 0.145157 |
| **46a** | **ER** | **appear vs claw** | **-86.41509** | **176.8302** | **-82.12116** | **172.2423** | **8.58786** | **0.0136512** |
| 46b | SYM | appear vs claw | -86.41509 | 176.8302 | -82.12116 | 172.2423 | 8.58786 | 0.0136512 |
| 46c | ARD | appear vs claw | -78.05674 | 164.1135 | -74.84498 | 165.6900 | 6.42351 | 0.169674 |
| **47a** | **ER** | **appear vs labellum** | **-86.55051** | **177.1010** | **-83.38377** | **174.7675** | **6.33348** | **0.0421408** |
| 47b | SYM | appear vs labellum | -86.55051 | 177.1010 | -83.38377 | 174.7675 | 6.33348 | 0.0421408 |
| 47c | ARD | appear vs labellum | -77.81554 | 163.6311 | -74.20754 | 164.4151 | 7.21601 | 0.124904 |
| 48a | ER | appear vs blotch | -87.53444 | 179.0689 | -86.04546 | 180.0909 | 2.97796 | 0.225602 |
| 48b | SYM | appear vs blotch | -87.53444 | 179.0689 | -86.04546 | 180.0909 | 2.97796 | 0.225602 |
| 48c | ARD | appear vs blotch | -82.89058 | 173.7812 | -78.95895 | 173.9179 | 7.86327 | 0.0967171 |
| **49a** | **ER** | **appear vs stigma** | **-82.95788** | **169.9158** | **-76.60013** | **161.2003** | **12.7155** | **0.00173326** |
| 49b | SYM | appear vs stigma | -82.95788 | 169.9158 | -76.60013 | 161.2003 | 12.7155 | 0.00173326 |
| 49c | ARD | appear vs stigma | -72.74475 | 153.4895 | -69.19565 | 154.3913 | 7.09821 | 0.130789 |
| 50a | ER | appear vs coiling | -53.82939 | 111.6588 | -53.31286 | 114.6257 | 1.03307 | 0.596584 |
| 50b | SYM | appear vs coiling | -53.82939 | 111.6588 | -53.31286 | 114.6257 | 1.03307 | 0.596584 |
| 50c | ARD | appear vs coiling | -49.14968 | 106.2994 | -47.35388 | 110.7078 | 3.59159 | 0.464089 |
| 51a | ER | appear vs calyx | -74.01693 | 152.0339 | -72.99576 | 153.9915 | 2.04234 | 0.360174 |
| 51b | SYM | appear vs calyx | -74.01693 | 152.0339 | -72.99576 | 153.9915 | 2.04234 | 0.360174 |
| 51c | ARD | appear vs calyx | -68.81802 | 145.6360 | -67.40067 | 150.8013 | 2.8347 | 0.585858 |
| 52a | ER | rachis vs bract | -60.86954 | 125.7391 | -60.86954 | 129.7391 | 0 | 1 |
| 52b | SYM | rachis vs bract | -60.86954 | 125.7391 | -60.86954 | 129.7391 | 0 | 1 |
| 52c | ARD | rachis vs bract | -60.86954 | 129.7391 | -60.86954 | 137.7391 | 9.53122e-10 | 1 |
| 53a | ER | rachis vs density | -78.70269 | 161.4054 | -76.80722 | 161.6144 | 3.79094 | 0.150248 |
| 53b | SYM | rachis vs density | -78.70269 | 161.4054 | -76.80722 | 161.6144 | 3.79094 | 0.150248 |
| 53c | ARD | rachis vs density | -77.53477 | 163.0695 | -73.52116 | 163.0423 | 8.02722 | 0.0905863 |
| 54a | ER | rachis vs cincinnus | -67.88974 | 139.7795 | -66.69282 | 141.3856 | 2.39383 | 2.39383 |
| 54b | SYM | rachis vs cincinnus | -67.88974 | 139.7795 | -66.69282 | 141.3856 | 2.39383 | 2.39383 |
| 54c | ARD | rachis vs cincinnus | -67.57153 | 143.1431 | -63.96367 | 143.9273 | 7.21572 | 0.124918 |
| 55a | ER | rachis vs flowers | -93.20786 | 190.4157 | -91.31995 | 190.6399 | 3.77582 | 0.151388 |
| 55b | SYM | rachis vs flowers | -93.20786 | 190.4157 | -91.31995 | 190.6399 | 3.77582 | 0.151388 |
| **55c** | **ARD** | **rachis vs flowers** | **-92.81454** | **193.6291** | **-84.60897** | **185.2179** | **16.4112** | **0.00251429** |
| **56a** | **ER** | **rachis vs claw** | **-92.07513** | **188.1503** | **-86.07896 1** | **80.1579** | **11.9923** | **0.00248826** |
| 56b | SYM | rachis vs claw | -92.07513 | 188.1503 | -86.07896 1 | 80.1579 | 11.9923 | 0.00248826 |
| 56c | ARD | rachis vs claw | -87.84937 | 183.6987 | -79.33518 | 174.6703 | 17.0284 | 0.00190854 |
| **57a** | **ER** | **rachis vs labellum** | **-92.21054** | **188.4211** | **-89.20447** | **186.4089** | **6.01213** | **0.0494859** |
| 57b | SYM | rachis vs labellum | -92.21054 | 188.4211 | -89.20447 | 186.4089 | 6.01213 | 0.0494859 |
| 57c | ARD | rachis vs labellum | -87.60818 | 183.2164 | -83.38125 | 182.7625 | 8.45386 | 0.0762981 |
| 58a | ER | rachis vs blotch | -93.19447 | 190.3889 | -92.11259 | 192.2252 | 2.16376 | 0.338957 |
| 58b | SYM | rachis vs blotch | -93.19447 | 190.3889 | -92.11259 | 192.2252 | 2.16376 | 0.338957 |
| 58c | ARD | rachis vs blotch | -92.68322 | 193.3664 | -89.20609 | 194.4122 | 6.95426 | 0.138325 |
| 59a | ER | rachis vs stigma | -88.61792 | 181.2358 | -77.76084 | 163.5217 | 21.7142 | 1.92678e-05 |
| 59b | SYM | rachis vs stigma | -88.61792 | 181.2358 | -77.76084 | 163.5217 | 21.7142 | 1.92678e-05 |
| **59c** | **ARD** | **rachis vs stigma** | **-82.53739** | **173.0748** | **-71.93187** | **159.8637** | **21.211** | **0.000287571** |
| 60a | ER | rachis vs coiling | -59.48945 | 122.9789 | -55.43943 | 118.8789 | 8.10003 | 0.0174221 |
| 60b | SYM | rachis vs coiling | -59.48945 | 122.9789 | -55.43943 | 118.8789 | 8.10003 | 0.0174221 |
| **60c** | **ARD** | **rachis vs coiling** | **-58.94232** | **125.8846** | **-50.22281** | **116.4456** | **17.439** | **0.00158785** |
| 61a | ER | rachis vs calyx | -79.67696 | 163.3539 | -77.82961 | 163.6592 | 3.69471 | 0.157654 |
| 61b | SYM | rachis vs calyx | -79.67696 | 163.3539 | -77.82961 | 163.6592 | 3.69471 | 0.157654 |
| 61c | ARD | rachis vs calyx | -78.61065 | 165.2213 | -74.82644 | 165.6529 | 7.56842 | 0.10873 |
| 62a | ER | calyx vs bract | -73.40381 | 150.8076 | -72.09182 | 152.1836 | 2.62397 | 0.269285 |
| 62b | SYM | calyx vs bract | -73.40381 | 150.8076 | -72.09182 | 152.1836 | 2.62397 | 0.269285 |
| 62c | ARD | calyx vs bract | -72.00879 | 152.0176 | -68.81895 | 153.6379 | 6.37969 | 0.172531 |
| 63a | ER | calyx vs density | -68.63523 | 141.2705 | -66.80716 | 141.6143 | 3.65612 | 0.160725 |
| 63b | SYM | calyx vs density | -68.63523 | 141.2705 | -66.80716 | 141.6143 | 3.65612 | 0.160725 |
| 63c | ARD | calyx vs density | -67.03441 | 142.0688 | -64.18768 | 144.3754 | 5.69344 | 0.223242 |
| 64a | ER | calyx vs cincinnus | -57.82227 | 119.6445 | -57.56405 | 123.1281 | 0.516442 | 0.772425 |
| 64b | SYM | calyx vs cincinnus | -57.82227 | 119.6445 | -57.56405 | 123.1281 | 0.516442 | 0.772425 |
| 64c | ARD | calyx vs cincinnus | -57.07117 | 122.1423 | -56.74871 | 129.4974 | 0.644915 | 0.957944 |
| 65a | ER | calyx vs flowers | -83.14039 | 170.2808 | -82.83421 | 173.6684 | 0.612368 | 0.736251 |
| 65b | SYM | calyx vs flowers | -83.14039 | 170.2808 | -82.83421 | 173.6684 | 0.612368 | 0.736251 |
| 65c | ARD | calyx vs flowers | -82.31418 | 172.6284 | -81.13355 | 178.2671 | 2.36125 | 0.66964 |
| 66a | ER | calyx vs claw | -82.00766 | 168.0153 | -80.81319 | 169.6264 | 2.38894 | 0.302864 |
| 66b | SYM | calyx vs claw | -82.00766 | 168.0153 | -80.81319 | 169.6264 | 2.38894 | 0.302864 |
| 66c | ARD | calyx vs claw | -77.34901 | 162.6980 | -75.83076 | 167.6615 | 3.03649 | 0.551738 |
| 67a | ER | calyx vs labellum | -82.14308 | 168.2862 | -77.69497 | 163.3899 | 8.8962 | 0.0117008 |
| 67b | SYM | calyx vs labellum | -82.14308 | 168.2862 | -77.69497 | 163.3899 | 8.8962 | 0.0117008 |
| **67c** | **ARD** | **calyx vs labellum** | **-77.10781** | **162.2156** | **-71.99931** | **159.9986** | **10.217** | **0.0369268** |
| 68a | ER | calyx vs blotch | -83.12701 | 170.2540 | -82.41920 | 172.8384 | 1.41561 | 0.492724 |
| 68b | SYM | calyx vs blotch | -83.12701 | 170.2540 | -82.41920 | 172.8384 | 1.41561 | 0.492724 |
| 68c | ARD | calyx vs blotch | -82.18286 | 172.3657 | -80.30480 | 176.6096 | 3.75611 | 0.440018 |
| 69a | ER | calyx vs stigma | -78.55045 | 161.1009 | -78.35643 | 164.7129 | 0.388046 | 0.823639 |
| 69b | SYM | calyx vs stigma | -78.55045 | 161.1009 | -78.35643 | 164.7129 | 0.388046 | 0.823639 |
| 69c | ARD | calyx vs stigma | -72.03703 | 152.0741 | -70.84055 | 157.6811 | 2.39295 | 0.663902 |
| 70a | ER | calyx vs coiling | -49.42199 | 102.8440 | -47.98615 | 103.9723 | 2.87168 | 0.237916 |
| 70b | SYM | calyx vs coiling | -49.42199 | 102.8440 | -47.98615 | 103.9723 | 2.87168 | 0.237916 |
| 70c | ARD | calyx vs coiling | -48.44195 | 104.8839 | -46.02617 | 108.0523 | 4.83157 | 0.305021 |
| 71a | ER | coiling vs bract | -53.21630 | 110.4326 | -52.45052 | 112.9010 | 1.53156 | 0.46497 |
| 71b | SYM | coiling vs bract | -53.21630 | 110.4326 | -52.45052 | 112.9010 | 1.53156 | 0.46497 |
| **71c** | **ARD** | **coiling vs bract** | **-52.34045** | **112.6809** | **-42.78478** | **101.5696** | **19.1113** | **0.000747325** |
| 72a | ER | coiling vs density | -48.44772 | 100.8954 | -46.97867 | 101.9573 | 2.9381 | 0.230145 |
| 72b | SYM | coiling vs density | -48.44772 | 100.8954 | -46.97867 | 101.9573 | 2.9381 | 0.230145 |
| 72c | ARD | coiling vs density | -47.36607 | 102.7321 | -44.48397 | 104.9679 | 5.7642 | 0.217464 |
| 73a | ER | coiling vs cincinnus | -37.63476 | 79.26952 | -35.20030 | 78.40060 | 4.86893 | 0.0876448 |
| 73b | SYM | coiling vs cincinnus | -37.63476 | 79.26952 | -35.20030 | 78.40060 | 4.86893 | 0.0876448 |
| 73c | ARD | coiling vs cincinnus | -37.40283 | 82.80566 | -34.50756 | 85.01511 | 5.79055 | 0.215346 |
| 74a | ER | coiling vs flowers | -62.95289 | 129.9058 | -60.16392 | 128.3278 | 5.57792 | 0.061485 |
| 74b | SYM | coiling vs flowers | -62.95289 | 129.9058 | -60.16392 | 128.3278 | 5.57792 | 0.061485 |
| 74c | ARD | coiling vs flowers | -62.64584 | 133.2917 | -58.37840 | 132.7568 | 8.53488 | 0.0738371 |
| 75a | ER | coiling vs claw | -61.82015 | 127.6403 | -60.73123 | 129.4625 | 2.17785 | 0.336578 |
| 75b | SYM | coiling vs claw | -61.82015 | 127.6403 | -60.73123 | 129.4625 | 2.17785 | 0.336578 |
| 75c | ARD | coiling vs claw | -57.68067 | 123.3613 | -56.78039 | 129.5608 | 1.80056 | 0.77238 |
| **76a** | **ER** | **coiling vs labellum** | **-61.95557** | **127.9111** | **-58.66436** | **125.3287** | **6.58242** | **0.0372089** |
| 76b | SYM | coiling vs labellum | -61.95557 | 127.9111 | -58.66436 | 125.3287 | 6.58242 | 0.0372089 |
| 76c | ARD | coiling vs labellum | -57.43948 | 122.879 | -56.27951 | 128.559 | 2.31993 | 0.677143 |
| 77a | ER | coiling vs blotch | -62.93950 | 129.8790 | -61.95137 | 131.9027 | 1.97626 | 0.372271 |
| 77b | SYM | coiling vs blotch | -62.93950 | 129.8790 | -61.95137 | 131.9027 | 1.97626 | 0.372271 |
| 77c | ARD | coiling vs blotch | -62.51452 | 133.0290 | -61.28434 | 138.5687 | 2.46036 | 0.651747 |
| 78a | ER | coiling vs stigma | -58.36294 | 120.7259 | -55.76392 | 119.5278 | 5.19804 | 0.0743465 |
| 78b | SYM | coiling vs stigma | -58.36294 | 120.7259 | -55.76392 | 119.5278 | 5.19804 | 0.0743465 |
| 78c | ARD | coiling vs stigma | -52.36869 | 112.7374 | -50.57349 | 117.1470 | 3.5904 | 0.464266 |

*character states (0 = white, 1 = non-white)

### **Appendix S1c.** Correlation matrix for discrete binary characters in *Hedychium*. (Likelihood Ratio/P-value; ER = Blue; ARD = Red)

|  | **slender** | **appear** | **rachis** | **bract** | **density** | **cincinnus** | **flowers** | **claw** | **Labellum**  **color** | **blotch** | **stigma** | **coiling** | **calyx** |
| --- | --- | --- | --- | --- | --- | --- | --- | --- | --- | --- | --- | --- | --- |
| **slender** |  | **-** | **-** | **-** | **-** | **-** | **-** | **-** | **-** | **-** | **-** | **-** | **11.3341/0.00345803** |
| **appear** |  |  | **29.2979/6.80019e-06** | **22.2066/0.000182308** | **-** | **-** | **-** | 8.58786/0.0136512 | **6.33348/0.0421408** | **-** | **12.7155/0.00173326** | **-** | **-** |
| **rachis** |  |  |  | **-** | **-** | **-** | **16.4112/0.00251429** | 11.9923/0.00248826 | **6.01213/0.0494859** | **-** | **21.211/0.000287571** | **17.439/0.00158785** | **-** |
| **bract** |  |  |  |  | **-** | **6.55631/0.0376978** | **13.9217/0.000948282** | 8.37223/0.0152053 | **6.37921/0.0411881** | **-** | **12.9107/0.0117205** | **19.1113/0.000747325** | **-** |
| **density** |  |  |  |  |  | **-** | **-** | **-** | **12.3889/0.00204071** | **-** | **8.4916/0.0143243** | **-** | **-** |
| **cincinnus** |  |  |  |  |  |  | **-** | **-** | **12.1917/0.00225221** | **7.06508/0.0292306** | **-** | **-** | **-** |
| **flowers** |  |  |  |  |  |  |  | **15.6299/0.000403659** | **-** | **-** | **16.6085/0.00230241** | **-** | **-** |
| **claw** |  |  |  |  |  |  |  |  | **-** | **-** | **38.5125/8.78366e-08** | **-** | **-** |
| **Labellum color** |  |  |  |  |  |  |  |  |  | **11.3357/0.0230393** | **9.64253/0.0468994** | **6.58242/0.0372089** | **10.217/0.0369268** |
| **blotch** |  |  |  |  |  |  |  |  |  |  | **-** | **-** | **-** |
| **stigma** |  |  |  |  |  |  |  |  |  |  |  | - | **-** |
| **coiling** |  |  |  |  |  |  |  |  |  |  |  |  | **-** |
| **calyx** |  |  |  |  |  |  |  |  |  |  |  |  |  |

### **Appendix S1d.** Estimation of phylogenetic signal for the discrete binary characters in *Hedychium* using Fritz-Purvis’D (Fritz and Purvis, 2010) and δ-statistic (Borges et al., 2019).

| **Sl. no.** | **Code** | **Character** | **Estimated D** | **Probability of E(D) resulting from no (random) phylogenetic structure** | **Probability of E(D) resulting from Brownian phylogenetic structure** | **δ-statistic** | ***P*-value** |
| --- | --- | --- | --- | --- | --- | --- | --- |
| **1** | **slender** | **Slender cylindrical inflorescence** | **-0.90** | **0.001** | **0.96** | **5.31** | **0** |
| 2 | appear | Inflorescence appears wider than long | 0.58 | 0.04 | 0.08 | *0.82* | *0.86* |
| 3 | rachis | Rachis exposure | 0.27 | 0 | 0.21 | **1.58** | **0.01** |
| 4 | bract | Inflorescence bract type | 0.24 | 0 | 0.26 | **3.61** | **0** |
| 5 | density | Floral density | 0.61 | 0.07 | 0.07 | **1.83** | **0** |
| **6** | **cincinnus** | **Cincinnus capacity** | **-0.73** | **0** | **0.98** | **37.35** | **0** |
| 7 | flowers | Flowers open per day | 0.78 | 0.14 | 0.003 | *0.09* | *0.38* |
| 8 | calyx | Relative length of bract and calyx | -0.03 | 0 | 0.55 | **3.98** | **0** |
| **9** | **coiling** | **Floral tube coiling** | **-1.03** | **0** | **0.99** | **91.8** | **0** |
| 10 | claw | Labellum claw | *0.71* | *0.08* | *0.02* | *0.45* | *0.68* |
| 11 | labellum | Labellum color* | 0.25 | 0 | 0.23 | **3.48** | **0** |
| 12 | blotch | Labellum blotch color | 0.93 | 0.33 | 0.002 | *0.26* | *0.14* |
| 13 | stigma | Stigma position | 0.80 | 0.19 | 0.01 | *0.66* | *0.91* |

*character states (0 = white, 1 = non-white)

### **Appendix S1e.** Estimation of Pagel’s λ and Blomberg’s K and the fit of different models of eight continuously variable floral characters in *Hedychium*.

| **Continuous character** | **Pagel’s λ** | **P-value (based on LR test)** | **Blomberg’s K** | **P-value (based on 1000 randomizations)** | **LnL/AIC/AICc** | | | | **α value** | **Sig^2^**  **^BM^** | **Sig^2^**  **^OU^** | **Sig^2^**  **^EB^** | **Sig^2^**  **^WN^** | **pchisq** | | |
| --- | --- | --- | --- | --- | --- | --- | --- | --- | --- | --- | --- | --- | --- | --- | --- | --- |
|  |  |  |  |  |  |  |  |  |  |  |  |  |  | **OU ~ BM** | **OU ~ EB** | **OU ~ WN** |
|  |  |  |  |  | **BM**  **(k=2)** | **OU**  **(k=3)** | **EB**  **(k=3)** | **WN**  **(k=2)** |  |  |  |  |  |  |  |  |
| Labellum length (LL) | 0.560955 | 0.0497094 | 0.277935 | 0.013 | -124.491113/252.982226/253.161330 | **-115.737985/237.475969/237.839606** | -124.491138/254.982275/255.345912 | -118.145765/240.291530/240.470634 | 0.60 | 0.797661 | 2.188738 | 0.797669 | 1.712092 | 2.863639e-05 | 2.863563e-05 | 0.02820391 |
| Labellum width (LW) | 0.37325 | 0.00271834 | 0.287244 | 0.01 | -131.598301/267.196602/267.375707 | **-116.970900/239.941800/240.305437** | -131.598333/269.196666/269.560303 | -119.369142/242.738285/242.917389 | 0.88 | 0.977254 | 3.137617 | 0.977264 | 1.772994 | 6.345877e-08 | 6.345667e-08 | 0.02851788 |
| Labellum notch depth (LND) | 0.154272 | 1 | 0.254497 | 0.031 | -78.610843/161.221687/161.400791 | **-69.181688/144.363376/144.727012** | -78.610867/163.221735/163.585371 | -71.149623/146.299246/146.478351 | 0.61 | 0.215039 | 0.579250 | 0.215041 | 0.447074 | 1.407964e-05 | 1.407928e-05 | 0.04726669 |
| Lateral staminode length (LSL) | 0.727388 | 0.00348259 | 0.341621 | 0.008 | -116.732057/237.464114/237.643219 | -**110.943861/227.887722/228.251358** | -116.732079/239.464158/239.827794 | -114.456213/232.912425/233.091530 | 0.34 | 0.639059 | 1.253059 | 0.639065 | 1.540798 | 0.0006679442 | 0.0006679284 | 0.008039276 |
| Lateral staminode width (LSW) | 0.0912012 | 0.498519 | 0.196952 | 0.329 | -85.389477/174.778954/174.958059 | -59.548610/125.097220/125.460857 | -85.389514/176.779028/177.142664 | **-60.442815/124.885630/125.064735** | 1.72 | 0.260993 | 1.132184 | 0.260996 | 0.329249 | 6.526714e-13 | 6.526468e-13 | 0.18112 |
| Floral tube length (FTL) | 0.665983 | 0.00120541 | 0.342443 | 0.006 | -172.461894/348.923788/349.102893 | **-167.129430/340.258860/340.622496** | -172.461915/350.923831/351.287467 | -173.508054/351.016107/351.195212 | 0.27 | 3.140941 | 5.495557 | 3.140972 | 8.326939 | 0.001091861 | 0.001091836 | 0.0003546323 |
| Filament length (FL) | 0.252843 | 0.159547 | 0.243909 | 0.03 | -158.519741/321.039482/321.218586 | -138.476001/282.952002/283.315638 | -158.519774/323.039549/323.403185 | **-138.522054/281.044108/281.223213** | 2.72 | 2.108918 | 16.788786 | 2.108940 | 3.064535 | 2.428404e-10 | 2.428322e-10 | 0.7615167 |
| Anther length (AL) | 0.724888 | 0.000142726 | 0.398837 | 0.001 | -22.584395/49.168791/49.347895 | **-16.628369/39.256738/39.620375** | -22.584418/51.168837/51.532473 | -22.220755/48.441510/48.620614 | 0.32 | 0.043383 | 0.081629 | 0.043383 | 0.110471 | 0.0005577193 | 0.0005577055 | 0.0008247139 |

### **Appendix S1f.** Summary of parameters of PGLS based on log-transformed values with different models focusing on two characters.

| **Sl. no.** | **Model** | **Intercept** | **Slope** | **Lambda (ML)**  **(95.0% CI)** | **Residual standard error**  **(RSE)** | **Adjusted R^2^** | **F-statistic** | **P-value** |
| --- | --- | --- | --- | --- | --- | --- | --- | --- |
| 1 | LL vs LW | 1.859365 | 0.706101 | 0.536 (0.114, 0.855) | 0.3411 on 68 degrees of freedom | 0.478 | 64.18 on 1 and 68 DF | 2.093e-11 |
| 2 | LL vs LND | 1.67936 | 1.10563 | 0.652 (0.148, 0.901) | 0.4132 on 68 degrees of freedom | 0.3359 | 35.9 on 1 and 68 DF | 8.79e-08 |
| 3 | LL vs LSL | -0.085789 | 0.972408 | 0.000 (NA, 0.339) | 0.1574 on 68 degrees of freedom | 0.8488 | 388.3 on 1 and 68 DF | < 2.2e-16 |
| 4 | LL vs LSW | 2.41549 | 1.22171 | 0.758 (0.356, 0.945) | 0.4483 on 68 degrees of freedom | 0.3428 | 36.98 on 1 and 68 DF | 6.125e-08 |
| 5 | LL vs FTL | 1.517072 | 0.284896 | 0.395 (NA, 0.831) | 0.3554 on 68 degrees of freedom | 0.355 | 38.98 on 1 and 68 DF | 3.177e-08 |
| 6 | LL vs FL | 2.472872 | 0.178447 | 0.408 (NA, 0.815) | 0.4349 on 68 degrees of freedom | 0.04475 | 4.232 on 1 and 68 DF | 0.0435 |
| 7 | LL vs AL | 1.04575 | 2.12093 | 0.491 (NA, 0.812) | 0.4013 on 68 degrees of freedom | 0.2442 | 23.3 on 1 and 68 DF | 8.199e-06 |
| 8 | LW vs LND | 0.88925 | 0.67602 | 0.415 (0.131, 0.736) | 0.41 on 68 degrees of freedom | 0.1195 | 10.36 on 1 and 68 DF | 0.001973 |
| 9 | LW vs LSL | -0.32721 | 0.64686 | 0.404 (0.118, 0.736) | 0.3457 on 68 degrees of freedom | 0.3674 | 41.08 on 1 and 68 DF | 1.615e-08 |
| 10 | LW vs LSW | 0.79301 | 1.79078 | 0.594 (0.274, 0.843) | 0.2573 on 68 degrees of freedom | 0.7176 | 176.4 on 1 and 68 DF | < 2.2e-16 |
| 11 | LW vs FTL | 0.397513 | 0.240859 | 0.403 (0.126, 0.728) | 0.3734 on 68 degrees of freedom | 0.2611 | 25.38 on 1 and 68 DF | 3.694e-06 |
| 12 | LW vs FL | 1.942606 | -0.074789 | 0.424 (0.113, 0.763) | 0.4395 on 68 degrees of freedom | -0.003798 | 0.7389 on 1 and 68 DF | 0.393 |
| 13 | LW vs AL | 0.28085 | 1.50416 | 0.374 (0.064, 0.713) | 0.4016 on 68 degrees of freedom | 0.1242 | 10.79 on 1 and 68 DF | 0.001617 |
| 14 | LND vs LSL | 0.276240 | 0.272695 | 0.000 (NA, 0.621) | 0.1797 on 68 degrees of freedom | 0.2453 | 23.43 on 1 and 68 DF | 7.782e-06 |
| 15 | LND vs LSW | 0.99472 | 0.39703 | 0.320 (NA, 0.903) | 0.2135 on 68 degrees of freedom | 0.1044 | 9.043 on 1 and 68 DF | 0.003693 |
| 16 | LND vs FTL | 0.660624 | 0.091247 | 0.000 (NA, 0.405) | 0.1916 on 68 degrees of freedom | 0.1427 | 12.48 on 1 and 68 DF | 0.0007441 |
| 17 | LND vs FL | 0.660624 | 0.091247 | 0.000 (NA, 0.405) | 0.1916 on 68 degrees of freedom | 0.1427 | 12.48 on 1 and 68 DF | 0.0007441 |
| 18 | LND vs AL | 0.660624 | 0.091247 | 0.000 (NA, 0.405) | 0.1916 on 68 degrees of freedom | 0.1427 | 12.48 on 1 and 68 DF | 0.0007441 |
| 19 | LSL vs LSW | 2.53263 | 0.97803 | 0.837 (0.495, 0.971) | 0.4653 on 68 degrees of freedom | 0.2768 | 27.41 on 1 and 68 DF | 1.734e-06 |
| 20 | LSL vs FTL | 1.505862 | 0.297126 | 0.483 (NA, 0.901) | 0.3148 on 68 degrees of freedom | 0.4522 | 57.96 on 1 and 68 DF | 1.104e-10 |
| 21 | LSL vs FL | 2.307336 | 0.236812 | 0.609 (0.118, 0.901) | 0.4265 on 68 degrees of freedom | 0.1129 | 9.781 on 1 and 68 DF | 0.002596 |
| 22 | LSL vs AL | 0.98816 | 2.22362 | 0.686 (0.270, 0.911) | 0.3938 on 68 degrees of freedom | 0.3159 | 32.86 on 1 and 68 DF | 2.484e-07 |
| 23 | LSW vs FTL | 0.078708 | 0.088845 | 0.174 (NA, 0.567) | 0.167 on 68 degrees of freedom | 0.1732 | 15.46 on 1 and 68 DF | 0.0002003 |
| 24 | LSW vs FL | 0.665855 | -0.015825 | 0.116 (NA, 0.522) | 0.1816 on 68 degrees of freedom | -0.01238 | 0.1564 on 1 and 68 DF | 0.6937 |
| 25 | LSW vs AL | 0.041238 | 0.649101 | 0.000 (NA, 0.424) | 0.1657 on 68 degrees of freedom | 0.1287 | 11.2 on 1 and 68 DF | 0.001338 |
| 26 | FTL vs FL | 5.630439 | -0.098845 | 0.687 (0.245, 0.958) | 1.098 on 68 degrees of freedom | -0.01038 | 0.2913 on 1 and 68 DF | 0.5912 |
| 27 | FTL vs AL | 0.22351 | 5.59224 | 0.776 (0.431, 0.968) | 0.9254 on 68 degrees of freedom | 0.3819 | 43.64 on 1 and 68 DF | 7.218e-09 |
| 28 | FL vs AL | 2.67334 | 1.27919 | 0.000 (NA, 0.655) | 0.5293 on 68 degrees of freedom | 0.04515 | 4.263 on 1 and 68 DF | 0.04278 |
| 29 | LL vs LW+LND+LSL+LSW+FTL+FL+AL | - | - | 0.000 (NA, 0.236) | 0.1325 on 62 degrees of freedom | 0.8929 | 83.2 on 7 and 62 DF | < 2.2e-16 |
| 30 | LW vs LL+LND+LSL+LSW+FTL+FL+AL | - | - | 0.468 (0.180, 0.760) | 0.2101 on 62 degrees of freedom | 0.7803 | 36.01 on 7 and 62 DF | < 2.2e-16 |
| 31 | LND vs LL+LW+LSL+LSW+FTL+FL+AL | - | - | 0.000 (NA, 0.764) | 0.1703 on 62 degrees of freedom | 0.3223 | 5.689 on 7 and 62 DF | 4.181e-05 |
| 32 | LSL vs LL+LW+LND+LSW+FTL+FL+AL | - | - | 0.000 (NA, 0.355) | 0.1174 on 62 degrees of freedom | 0.9066 | 96.71 on 7 and 62 DF | < 2.2e-16 |
| 33 | LSW vs LL+LW+LND+LSL+FTL+FL+AL | - | - | 0.305 (NA, 0.702) | 0.1086 on 62 degrees of freedom | 0.6845 | 22.39 on 7 and 62 DF | 8.82e-15 |
| 34 | FTL vs LL+LW+LND+LSL+LSW+FL+AL | - | - | 0.608 (0.189, 0.926) | 0.6571 on 63 degrees of freedom | 0.5976 | 18.08 on 6 and 63 DF | 4.521e-12 |
| 35 | FL vs LL+LW+LND+LSL+LSW+FTL+AL | - | - | 0.686 (0.267, 0.911) | 0.5477 on 63 degrees of freedom | 0.4223 | 9.408 on 6 and 63 DF | 2.307e-07 |
| 36 | AL vs LL+LW+LND+LSL+LSW+FTL+FL | - | - | 0.774 (0.481, 0.943) | 0.1027 on 63 degrees of freedom | 0.3876 | 8.278 on 6 and 63 DF | 1.276e-06 |

### **Appendix S1g.** Correlation matrix for continuous characters in *Hedychium* (Adjusted R^2^/P-value).

|  | **LL** | **LW** | **LND** | **LSL** | **LSW** | **FTL** | **FL** | **AL** |
| --- | --- | --- | --- | --- | --- | --- | --- | --- |
| **LL** |  | **0.478/2.093e-11** | **0.3359/8.79e-08** | **0.8488/< 2.2e-16** | **0.3428/6.125e-08** | **0.355/3.177e-08** | **0.04475/0.0435** | **0.2442/8.199e-06** |
| **LW** |  |  | **0.1195/0.001973** | **0.3674/1.615e-08** | **0.7176/< 2.2e-16** | **0.2611/3.694e-06** | **-0.003798/0.393** | **0.1242/0.001617** |
| **LND** |  |  |  | **0.2453/7.782e-06** | **0.1044/0.003693** | **0.1427/0.0007441** | **0.1427/0.0007441** | **0.1427/0.0007441** |
| **LSL** |  |  |  |  | **0.2768/1.734e-06** | **0.4522/1.104e-10** | **0.1129/0.002596** | **0.3159/2.484e-07** |
| **LSW** |  |  |  |  |  | **0.1732/0.0002003** | -0.01238/0.6937 | **0.1287/0.001338** |
| **FTL** |  |  |  |  |  |  | -0.01038/0.5912 | **0.3819/7.218e-09** |
| **FL** |  |  |  |  |  |  |  | **0.04515/0.04278** |
| **AL** |  |  |  |  |  |  |  |  |

### **Appendix S1h.** Correlation matrix for continuous characters in *Hedychium* (Adjusted R^2^/P-value).

|  | **LW+LND+LSL+LSW+FTL+FL+AL** | **LL+LND+LSL+LSW+FTL+FL+AL** | **LL+LW+LSL+LSW+FTL+FL+AL** | **LL+LW+LND+LSW+FTL+FL+AL** | **LL+LW+LND+LSL+FTL+FL+AL** | **LL+LW+LND+LSL+LSW+FL+AL** | **LL+LW+LND+LSL+LSW+FTL+AL** | **LL+LW+LND+LSL+LSW+FTL+FL** |
| --- | --- | --- | --- | --- | --- | --- | --- | --- |
| **LL** | **0.8929/< 2.2e-16** | **1** | **2** | **3** | **4** | **5** | **6** | **7** |
| **LW** |  | **0.7803/< 2.2e-16** | **8** | **9** | **10** | **11** | **12** | **13** |
| **LND** |  |  | **0.3223/4.181e-05** | **14** | **15** | **16** | **17** | **18** |
| **LSL** |  |  |  | **0.9066 /< 2.2e-16** | **19** | **20** | **21** | **22** |
| **LSW** |  |  |  |  | **0.6845/8.82e-15** | **23** | **24** | **25** |
| **FTL** |  |  |  |  |  | **0.5976/4.521e-12** | **26** | **27** |
| **FL** |  |  |  |  |  |  | **0.4223/2.307e-07** | **28** |
| **AL** |  |  |  |  |  |  |  | **0.3876/1.276e-06** |

### **Appendix S1i.** Evolutionary transitions for the discrete characters in *Hedychium*.

#### ***See attached excel sheet.***
